# The GATA3 X308_Splice breast cancer mutation is a hormone context-dependent oncogenic driver

**DOI:** 10.1101/664367

**Authors:** Natascha Hruschka, Maria Subijana, Osvaldo Graña-Castro, Francisco Del Cano-Ochoa, Laia Paré Brunet, Ana Sagrera, Aurelien De Reynies, David Andreu, Joe Sutton, Igor Chernukhin, Suet-Feung Chin, Carlos Caldas, Ana Lluch, Octavio Burgués, Begoña Bermejo, Santiago Ramón-Maiques, Jason S Carroll, Aleix Prat, Francisco X Real, Paola Martinelli

## Abstract

As the catalogue of oncogenic driver mutations is expanding, it is becoming clear that alterations in a given gene should not be lumped into one single class, since they might have different functions. The transcription factor *GATA3* is a paradigm of this. Here, we address the functions of the most common *GATA3* mutation (X308_Splice) which generates a neoprotein that we designate as neoGATA3, associated with good patient prognosis. Based on extensive analyses of molecular and clinical data from approximately 3000 breast cancer patients, supported by mechanistic studies *in vitro*, we show that neoGATA3 interferes with the transcriptional programs controlled by estrogen and progesterone receptors, without fully abrogating them. This has opposite outputs in the pre- or post-menopausal hormonal context, having pro- or anti-proliferative effects, respectively. NeoGATA3 is an example of a context- and stage-dependent driver mutation. Our data call for functional analyses of putative cancer drivers to guide clinical application.

## Introduction

The recent large scale genomics studies have produced an expanding catalogue of cancer-driving somatic mutations, which now needs to be translated into biological and clinically applicable knowledge [1]. One limitation of many studies of cancer drivers is the tendency to lump all the genetic alterations occurring in one gene into a single class, which can lead to inconclusive or confusing results when patients are stratified in a binary fashion according to the presence of alterations, as different genetic alterations might have distinct effects [2, 3, 4]. The *GATA3* transcription factor is emerging as a paradigm of a gene where multiple classes of mutations occur, having distinct biological and clinical output [5, 6, 7, 8]. In the case of *GATA3*, this seems to be rather specific for breast cancer (BC), where *GATA3* is mutated in around 11% of cases and shows a characteristic mutational pattern, which differs from other tumor types [2, 3].

Several evidences indicate that GATA3 is involved in the activation of the mammary differentiation program: 1) in normal tissue, it is necessary for the formation of the luminal compartment [9]; 2) GATA3 expression in BC strongly correlates with estrogen receptor (ER) expression [10]; 3) GATA3 functions in a complex with FOXA1 and ER to recruit RNA polymerase and enhance transcription of ER-responsive genes [11]; and 4) ectopic expression in GATA3-negative basal-like BC cells is sufficient to induce luminal differentiation and inhibit tumor dissemination [12]. Consistent with this function, GATA3 expression decreases during the progression to metastatic BC [13]. The high frequency of *GATA3* mutations in BC supports the idea that they are driver mutations, but whether they result in loss-of-function (LOF) or gain-of-function (GOF) is not fully clear. Most *GATA3* mutations are rare or unique frameshift indels (insertion/deletions) distributed along the 3’ end of the gene (Figure 1A), consistent with the classical mutational pattern of a tumor suppressor and therefore suggesting a LOF [2]. However, they are typically heterozygous and the expression of the wild type allele is retained [14]. A few mutations concentrate in two clusters in exon 5 and 6, including some “hotspots” or “warmspots”, supporting the idea that they might generate GOF, instead. The question on whether *GATA3* mutations are true oncogenic drivers is also still open: while some *in vitro* and *in vivo* data suggest that they might favor tumor growth [6, 8, 15], in general they are associated with longer survival [2] and better response to endocrine therapy [16]. A recent study identified four classes of frameshift mutations in *GATA3*, which were suggested to have distinct functions: 1) ZnFn2 mutations, occurring within the C-terminal Zn finger, required for specific binding to GATA motifs; 2) splice mutations, occurring mainly between intron 4 and exon 5; 3) truncating mutations, occurring downstream of the C-terminal Zn finger; and 4) extension mutations, occurring in exon 6 and disrupting the stop codon [6]. ZnFn2 mutations produce a highly stable truncated protein lacking the C-terminal Zn finger, showing low affinity for DNA and altered transcriptional activity, and are associated with poor outcome when compared with other *GATA3* mutations [6, 17]. On the other hand, extension mutations produce a longer protein that modulates the sensitivity to drugs [5]. The effect of the splice and truncating mutations remains unknown.

**Figure 1:**
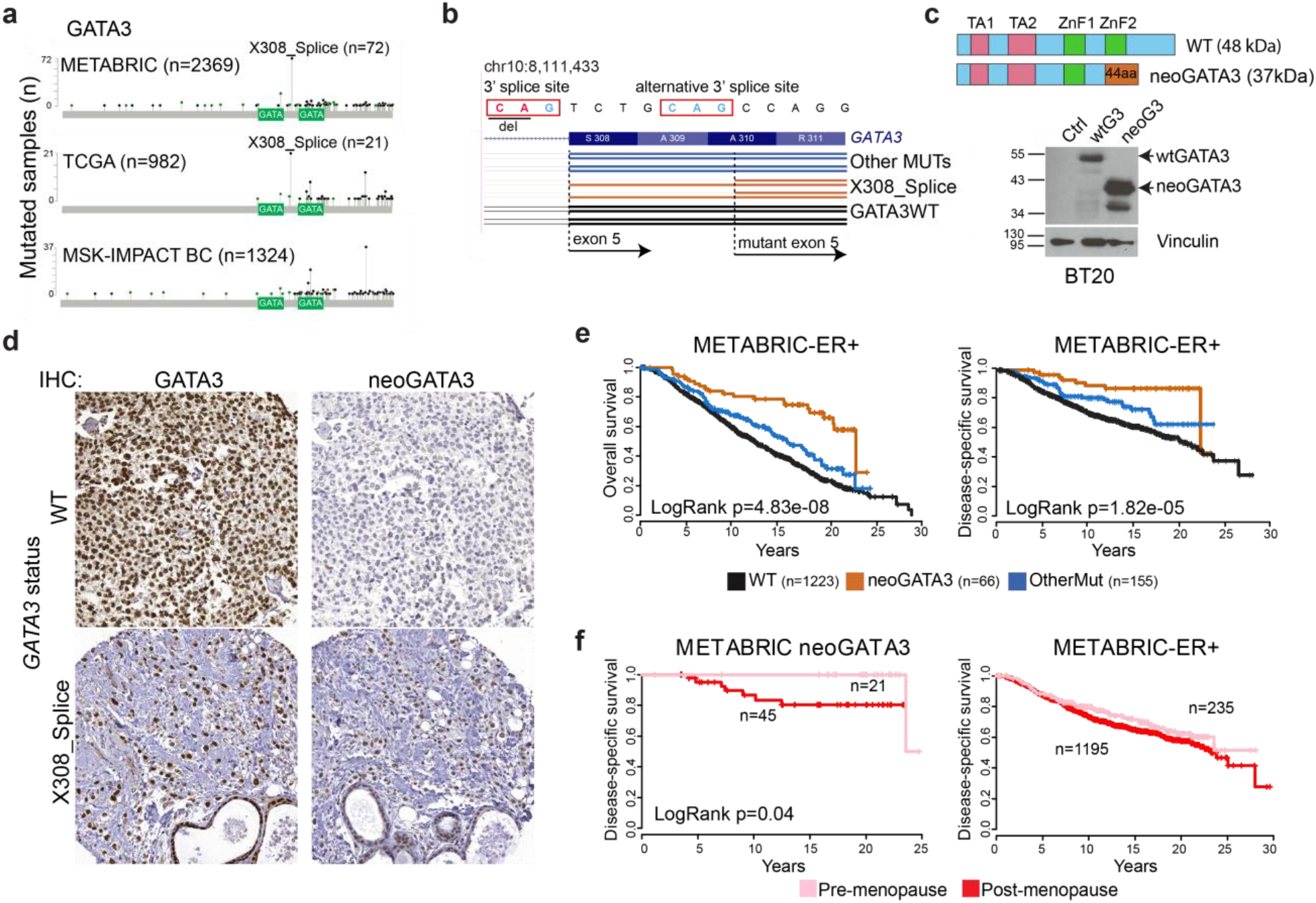
A hotspot splice-disrupting *GATA3* mutation correlates with good outcome in breast cancer. (**a**) Distribution of the *GATA3* mutations in the METABRIC, TCGA-BRCA, and MSK-IMPACT cohorts (only BC patients are shown for the latter), as displayed in the cBioPortal for Cancer Genomics webpage. The two GATA boxes indicate the two Zn finger DNA binding domains of the GATA3 protein. (**b**) UCSC genome browser-derived scheme of the mutant transcript identified in tumors carrying the X308_Splice mutation, compared with tumors with wt *GATA3* or with any other *GATA3* mutation. (**c**) Top: Schematic representation of wt GATA3, compared with the predicted neoGATA3 protein resulting from the X308_Splice mutation. Bottom: western blot showing the expression of a wild type GATA3 cDNA (wtG3) or the truncated transcript identified in the tumors with the X308_Splice mutation (neoGATA3). Black arrows indicate the proteins of the expected size. (**d**) Representative IHC images using either the N-ter GATA3 antibody - recognizing both wt and mutant GATA3 (left) - or the neoGATA3-specific antibody (right) on tumors carrying wild type GATA3 (top), or the X308_Splice mutation (bottom). (**e**) Kaplan-Meier survival curves of the METABRIC ER+ patients stratified according to *GATA3* status (WT= wild type, neoGATA3 = all mutations producing a neoGATA3-like peptide, OtherMut = all other mutations in GATA3). Left: overall survival, right: disease-specific survival. (**f**) Kaplan-Meier curves showing the disease-specific survival of the neoGATA3 (left) or all METABRIC ER+ (right) patients stratified according to the pre/post menopause status. Cox proportional hazard P value is indicated.

Here, we investigated the effects of the most prevalent *GATA3* hotspot somatic splice mutation (X308_Splice). This mutation, like 5 additional ones producing a partially or fully identical C-terminal peptide, correlates with significantly better outcome in patients and is associated with a specific gene expression signature, characterized by altered ER-dependent transcriptional program and reduced E2F target genes. The combined analysis of patient-derived data and *in vitro* experiments with breast cancer cell lines shows that the mutant protein - which we designate as “neoGATA3” interferes with the function of both ER and PR, blunting, without abrogating, their downstream programs. This has distinct biological outputs depending on the hormonal context: neoGATA3-expressing cells have a proliferative advantage when both estrogen and progesterone levels are high (before menopause) while they display a growth disadvantage when estrogen prevails (after menopause). Our data suggest the existence of stage-dependent oncogenic effects of driver mutations.

## Results

### The *GATA3* X308_splice mutation produces a unique neopeptide

The most common *GATA3* mutation reported in multiple BC genomic studies is a 2nt deletion in intron 4 disrupting the 3’ splice site (X308_Splice, Figure 1a). The predicted effect is a transcript lacking 7 nucleotides [7, 14] which we successfully identified in RNA-Seq data from 15/19 TCGA-BRCA samples carrying the X308_Splice mutation but not in 20/20 tumors with either wild type *GATA3* or carrying other mutations in the gene (Figure 1b). The mutant transcript was detected by RT-qPCR in 4/4 independent luminal A/B tumors carrying the X308_Splice mutation and in 0/7 without it (Supplementary Figure 1a). The loss of 7nt causes a frameshift, leading to a GATA3 protein - designated neoGATA3 - lacking residues 308-444, encompassing the second ZnFn, and containing instead a novel 44aa C-terminal sequence that does not display homology to any other human protein sequence (Figure 1c). Using western blotting, a polyclonal antiserum raised against the novel 44aa peptide specifically recognized a shorter GATA3 protein of the expected size (37KDa) exclusively in tumor cells carrying the mutation and in BC lacking endogenous GATA3 transduced with the mutant cDNA (Figure 1c and Supplementary Figure 1b). These antibodies allowed the detection of neoGATA3 in a tissue microarray containing 100 luminal A/B tumors with high sensitivity (90%) and specificity (94%) (Figure 1d).

The hotspot nature of the X308_Splice mutation suggests that it acts as an oncogenic driver and that other mutations might give rise to proteins with a similar C-terminal peptide. Indeed, we identified 5 additional mutations, detected in 6 METABRIC and 1 TCGA-BRCA samples, producing fully or partially identical C-terminal peptides. One of them is a 2nt insertion at codon Q321, which was found after re-sequencing one METABRIC sample (MB-0114), that showed immunoreactivity with the mutant-specific antibodies and had been originally genotyped as *GATA3*-wild type (Supplementary Figure 1c). In all, at least 6 different mutations, found in 78/2369 (3.3%) METABRIC samples and in 22/988 (2.2%) TCGA-BRCA samples, produce a neoGATA3-like peptide (Supplementary Table 1).

### Patients with neoGATA3-mutant tumors display good prognosis

To understand the clinical significance of neoGATA3 mutations, we analyzed the METABRIC cohort, where clinical data are available for 1673 patients, 231 (13.8%) of whom correspond to *GATA3*-mutant tumors. Among the latter, 66 (28.6%) had neoGATA3-type mutations and 165 (71.4%) had other mutations (OtherMut). NeoGATA3 mutations were significantly associated with lower tumor stage, grade, and size and with expression of progesterone receptor (PR) (Supplementary Figure 2a-d), all factors predicting better outcome. Consistently, patients with neoGATA3-mutant tumors had a significantly better overall survival (OS) and, most importantly, disease-specific survival (DSS) compared to both patients with wild type *GATA3* (WT) and those carrying any other *GATA3* mutation (OtherMut) (OS: LogRank P=7.58e-08, DSS: LogRank P=7.64e-07, Supplementary Figure 3a).

Most *GATA3* mutations were found in ER+ tumors; in particular, neoGATA3 mutations were exclusive for patients with ER+ tumors (Supplementary Figure 3b), which have better outcome [3, 18]. We therefore limited our analyses to these patients. The presence of neoGATA3 mutations was again strongly associated with significantly longer OS and DSS in this patient subgroup (OS: LogRank P=4.83e-08, DSS: LogRank P=1.82e-05, Figure 1e). A similar tendency towards longer DSS and disease-free survival (DFS) was observed for patients with neoGATA3-mutant tumors in the TCGA-BRCA ER+ cohort although the differences were not statistically significant, likely due to the smaller sample size (Supplementary Figure 3b). Univariate and multivariate analyses showed that neoGATA3 is an independent prognostic factor of longer OS and DSS in the METABRIC cohort (OS: HR=0.58, P=0.02; DSS: HR=0.46, P=0.034, Supplementary Tables 2-3). Consistent with the association of neoGATA3 mutations with a better prognosis, only 1/1324 (0.08%) patients with metastatic breast cancer harbored this genetic alteration [19] indicating that tumors with neoGATA3 mutations only metastasize exceptionally (P<0.0001, Figure 1a).

To get insight into the molecular features of tumors harboring neoGATA3 mutations, we derived a gene expression signature based on a training set composed of 981 TCGA-BRCA samples (19 neoGATA3 mutations). This signature could identify the neoGATA3 mutant tumors from the METABRIC series (n=2001 samples with expression and mutation data, 63 neoGATA3) with a sensitivity of 68.3% and specificity of 80.5% both when applied as a continuous variable and as binary classifier. When the signature was used to classify the samples from a cohort of patients with no available mutational data [20], patients with tumors classified as positive for the neoGATA3-signature (either as continuous or binary classifier) showed significantly longer DFS (LogRank P=0.004, not shown).

Strikingly, approximately one third of the neoGATA3 mutations occurred in pre-menopausal METABRIC patients (Supplementary Figure 3d, P=0.0004), who also had an extremely good prognosis when compared with the post-menopausal patients with neoGATA3 mutations (LogRank P=0.04) (Figure 1f, left). On the contrary, no difference between pre- and post-menopausal patients was observed in the whole METABRIC ER+ cohort (Figure 1g, right) or within patients with ER+ tumors having WT *GATA3* or other *GATA3* mutations (not shown). This suggested that the effect of the neoGATA3 mutations might be affected by age or age-related factors, including the hormonal context.

In summary, neoGATA3 mutations and the associated transcriptional signature are found in non-aggressive, non-metastatic, breast tumors of good prognosis.

### Tumors with neoGATA3 mutations show changes in the immune microenvironment, not consistent with a T-cell mediated acute immune response

A recent study identified the C-terminal neopeptide of neoGATA3 as a potential neoantigen and suggested that it might induce an anti-tumor T-cell-dependent immune response and the activation of immune checkpoints [21]. To assess the immune landscape of neoGATA3-mutant tumors, we used MCP-counter to deconvolute the expression of immune markers and estimate the abundance of different cell populations [22]. A significant decrease in “T cell” (P=0.002), “CD8+ T cell” (P=8.12e-06), “NK cell” (P=8.1e-05), and “Cytotoxic lymphocyte” (P=2.9e-04) signatures was observed in the neoGATA3 tumors of the METABRIC series, when compared with the WT tumors (Figure 2a). These observations were confirmed at the single gene level: the expression of the T-cell markers CD8A (P=0.015) and CD8B (P=1.23e-06), and of the immune checkpoint proteins PD-1 (P=0.005) and PD-L1 (P=0.033) was lower in neoGATA3 compared with WT tumors among the METABRIC patients (Figure 2b). We then analyzed the amount of CD8+ T cells in a set of FFPE sections from WT (n=6) and neoGATA3 (n=9) tumors by IHC for CD8α protein. In accordance to the gene expression data, CD8+ cells were significantly less abundant in neoGATA3 tumors (Figure 2c, P=0.042). No significant differences at the single gene level were observed in the neoGATA3 tumors of the TCGA cohort (Supplementary Figure 4a) but the “C4-lymphocyte depleted” immunoscore [23] was over-represented among the neoGATA3 tumors (4/17 neoGATA3 versus 47/525 WT), although statistical significance was not reached (P=0.06, not shown).

**Figure 2:**
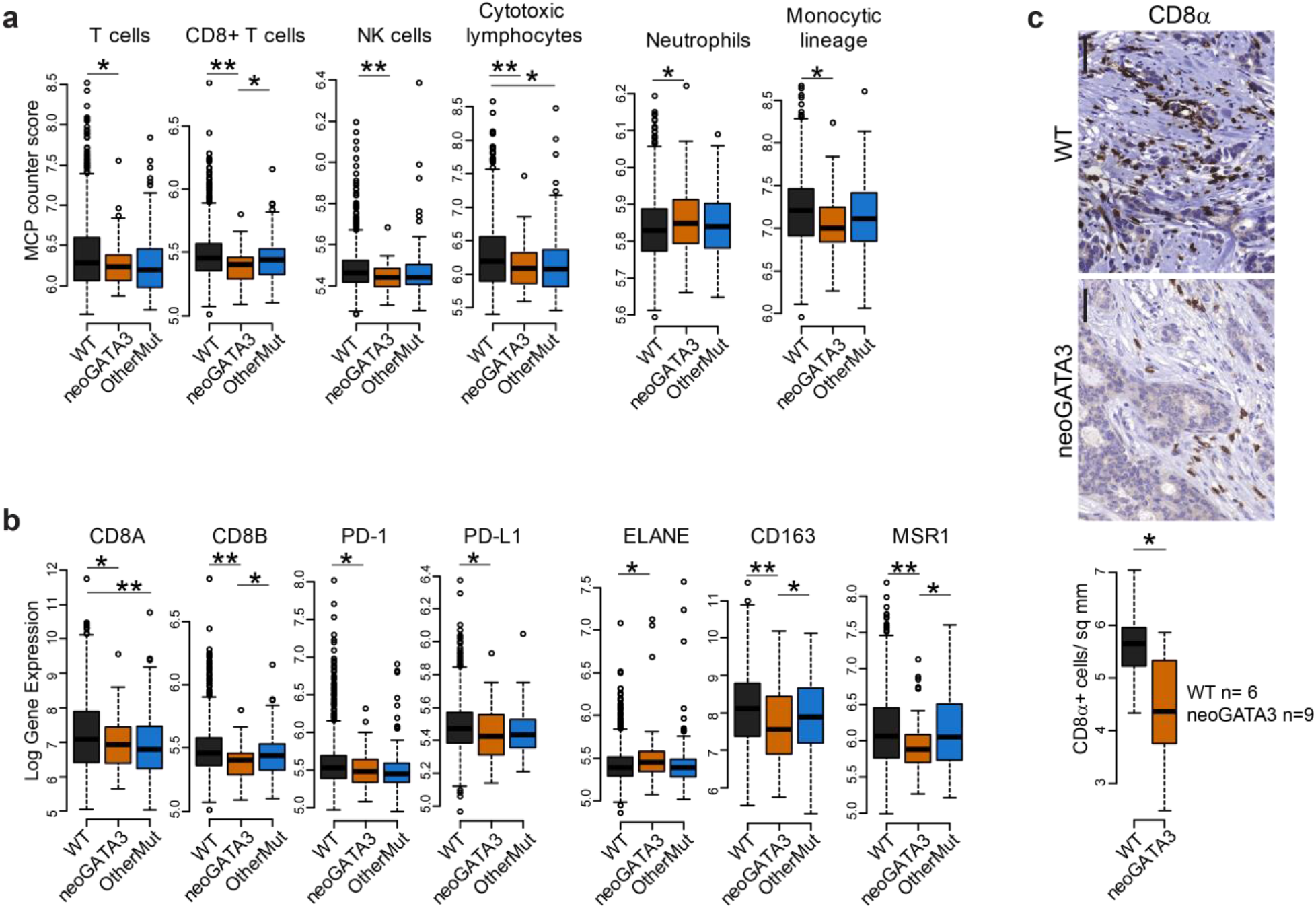
NeoGATA3-mutant tumors do not display a prominent immune response. (**a**) MCP counter scores for the indicated immune cell populations in the three groups of METABRIC tumors (WT n=1205, neoGATA3 n=65, OtherMut n=161). (**b**) Gene expression levels in tumors of the METABRIC cohort, divided in the three groups according to *GATA3* status (WT n=1189, neoGATA3 n=66, OtherMut n=155). (**c**) Representative IHC images of CD8α-positive cells in one tumor with wt GATA3 and in one with neoGATA3. Quantification of the staining of WT (n=6) and neoGATA3 (n=9) tumors is shown under the microphotographs. Two-sided Student’s T test *P<0.05, **P<0.01.

The MCP counter analyses revealed that the “Neutrophil” signature was upregulated (P=0.034) and the “Monocytic lineage” signature was down-regulated (P=0.0015) in the neoGATA3 tumors compared with WT, suggesting a complex modulation of the immune landscape in tumors carrying neoGATA3 mutations (Figure 2a). Accordingly, the neutrophil marker ELANE (P=0.035) was significantly increased, whereas the M2-macrophage markers CD163 (P=0.0001) and MSR1 (coding for CD204, P=2.4e-04) were decreased in the neoGATA3 METABRIC tumors compared with WT (Figure 2b) and showed similar tendencies in the TCGA-BRCA samples (Supplementary Figure 4b). For some of the indicated markers, a significant difference was also observed when comparing neoGATA3 with OtherMut tumors, again supporting distinct functions of the different mutations (Figure 2a,b).

Altogether, these data indicate that the neoGATA3 tumors display a distinct immune composition which does not fit with an active T cell-dependent immune response.

### Tumors with neoGATA3 mutations show decreased cell cycle progression and altered ER- and PR-dependent programs

To acquire a broader view of the molecular features of neoGATA3-mutant ER+ tumors, we analyzed the transcriptomic data from the METABRIC cohort. GSEA of the genes differentially expressed in the neoGATA3 tumors, compared with all other tumors, revealed a strong down-regulation of cell cycle- and inflammation-related gene sets (Figure 3a). Accordingly, mRNA levels of several cyclins, as well as PCNA and MKI67, were found to be lower in the neoGATA3 METABRIC tumors (Supplementary Figure 5a) consistent with the better prognosis observed in patients carrying neoGATA3 mutations. Similar differences were observed at the protein level in the TCGA series with reverse-phase protein arrays (RPPA) (Figure 3b). To get insight into the mediators of the neoGATA3-associated cell-cycle features, we computed the enrichment of transcription factor binding motifs on the promoter of the genes differentially regulated in neoGATA3 tumors. The E2F motifs were strongly enriched among the down-regulated genes (Figure 3c) and E2F2 and E2F4 mRNAs were significantly reduced in the neoGATA3 tumors, both compared with the WT (E2F2 P=0.027, E2F4 P=0.024) and with OtherMut (E2F2 P=0.035, E2F4 P=0.025) tumors (Figure 3d). There was no enrichment of GATA binding motifs in these genes. These findings indicate that the transcriptome of tumors carrying neoGATA3 might result from the blunting of the E2F-dependent program, which appears to be GATA3-independent.

**Figure 3:**
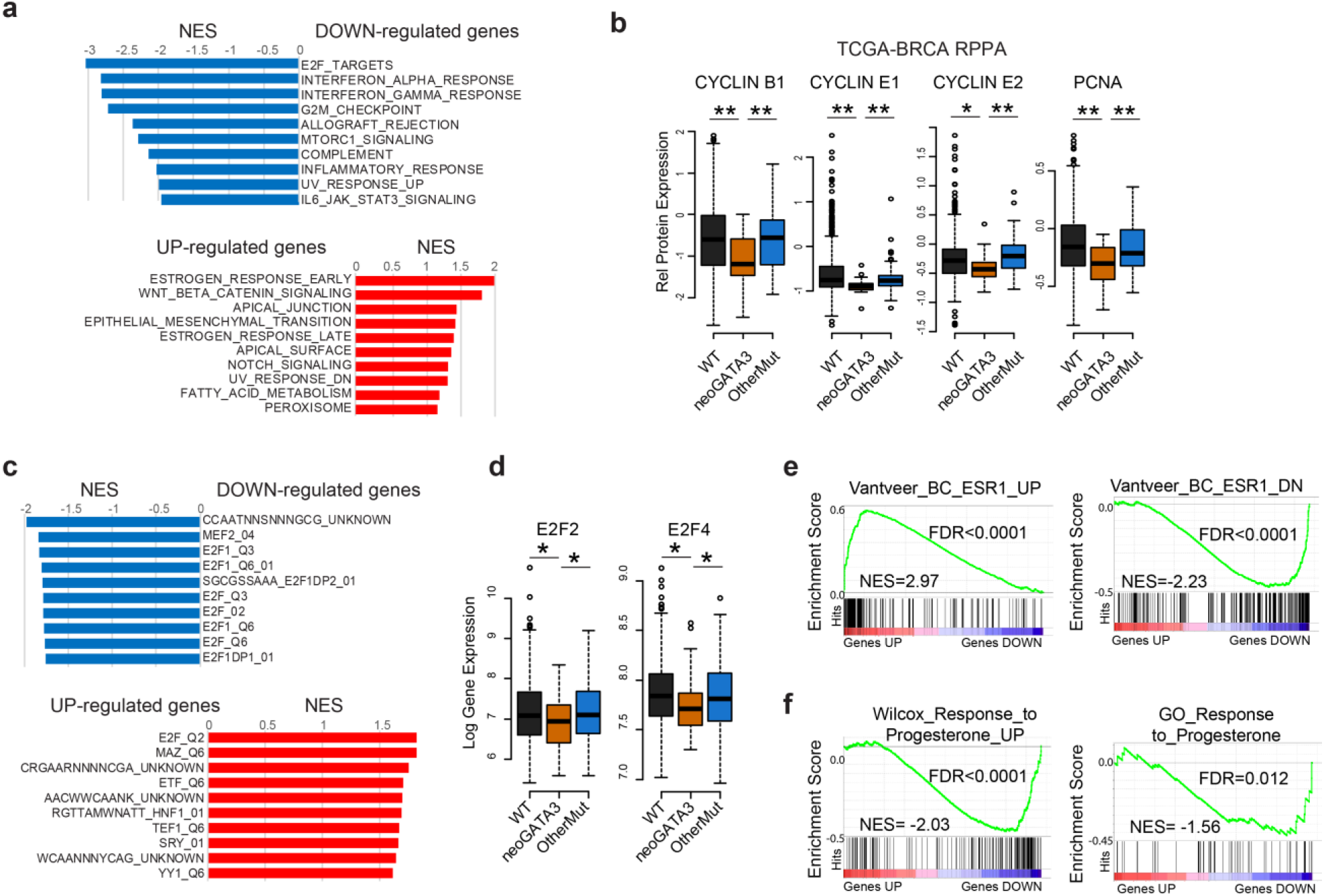
NeoGATA3 is associated with ER- and E2F-dependent transcriptional programs. (**a**) GSEA on the ranked list of genes differentially regulated in neoGATA3 tumors (n=65) compared with all other tumors of the METABRIC cohort (n=1345). The “Hallmarks” collection of gene sets was interrogated. The graphs show the normalized enrichment score (NES) of the 10 gene sets most significantly enriched among the up-regulated (red) and the down-regulated (blue) genes. FDR<0.05 for all gene sets shown. (**b**) RPPA data from the TCGA cohort showing expression levels of the indicated proteins in the three tumor groups (WT n=533, neoGATA3 n=18, OtherMut n=75). (**c**) GSEA as in (a) using the geneset collection “C3_tft” from the MsigDB. (**d**) E2F2 and E2F4 mRNA expression levels in the METABRIC tumors of the three groups (WT n=1189, neoGATA3 n=66, OtherMut n=155). Two-sided Student’s T-test *P<0.05. (**e**) Enrichment plots for the GSEA comparing the ranked list of genes regulated in the neoGATA3 tumors (n= (red-to-blue bar) with the indicated gene sets. The profile of the running enrichment score is shown in green, black bars represent the genes included in the gene set. FDR<0.05 for all gene sets shown. (**f**) Enrichment plots for the indicated gene sets within the genes up- or down-regulated in the METABRIC pre-menopause patients, comparing neoGATA3 versus all others.

Among the genes up-regulated in neoGATA3 tumors, we identified a significant enrichment of gene sets relative to the estrogen response (both early and late), WNT/β-catenin signaling, and the apical junctions (Figure 3a). Given the prominent enrichment of the “estrogen response” pathways, the known function of GATA3 as pioneer factor for ER genomic binding [11], and the exclusiveness of neoGATA3 mutations to ER+ tumors, we restricted the GSEA analyses to gene sets that were defined as up- or down-regulated in ER+ versus ER-tumors (Supplementary Table 3) and confirmed that the ER-associated transcriptome was significantly up-regulated in the neoGATA3 tumors both in the METABRIC and in the TCGA-BRCA cohort even within the ER+ tumors, suggesting a selective modulation of the ER program (Figure 3e, Supplementary Figure 5b). Furthermore, the neoGATA3-associated transcriptome showed a significant positive correlation with a gene signature of good prognosis and negative correlation with a signature of bad prognosis [24] (Supplementary Figure 5c), supporting a skewing of the ER-dependent program towards a less aggressive tumor phenotype. When analyzing the transcription factor binding motifs enriched in the up-regulated genes, we observed that the E2F motif was also the most enriched, indicating a striking functional interaction with this family of transcription factors (Figure 3c). DNA binding motifs for MAZ, ETF, HNF1, TEAD (TEF1), SRY, and YY1 were also significantly enriched, while the GATA motif was not significantly enriched, as observed for the down-regulated genes.

Recent work has shown that the ZnFn2 *GATA3* mutations interfere with the expression of the *PGR* gene, coding for PR [6]. Progesterone-related gene signatures were down-regulated in neoGATA3 tumors (Supplementary Figure 5d) even among pre-menopausal patients (Figure 3f). Interestingly, PGR was higher in neoGATA3 pre-menopausal tumors, compared to both WT (P=0.01) and OtherMut (P=0.006) (Supplementary Figure 5e), indicating that the two types of mutations exhibit distinct mechanisms of interference with PR activity [6].

Altogether, these data support that neoGATA3 modulates both the ER- and the PR-dependent programs in tumors, and the final major output is a general reduction of the E2F-driven proliferation.

### The neoGATA3 protein is more stable and shows altered DNA binding

To investigate the molecular mechanisms that could account for the association of neoGATA3 mutations with good prognosis, we searched for cellular models carrying the X308_Splice mutation. None of the 36 BC cell lines that we analyzed, which included 10 ER+ lines, harbored this mutation, suggesting that the growth of tumors with neoGATA3 mutations is not favored *in vitro*. We therefore relied on lentiviral-based transduction of BC cells with HA-tagged neoGATA3 cDNA (HA-neoG3) and used Flag-tagged wild type GATA3 cDNA (Flag-wtG3), as well as an empty vector as controls.

We first analyzed the general biochemical properties of neoGATA3 in BC cells lacking detectable endogenous GATA3. Expression of HA-neoG3 or Flag-wtG3 in BT20 (Figure 1c) and MDA-MB-468 cells (Supplementary Figure 6a), followed by cycloheximide treatment, revealed that neoGATA3 is markedly more stable than the wild type protein (estimated half-life >16h vs. 2h, respectively) (Figure 4a and Supplementary Figure 6b). Treatment with a proteasome inhibitor (MG132) increased the half-life of wtGATA3 but not neoGATA3, suggesting that the mutant protein is not affected by proteasome activity (Supplementary Figure 6c). Of note, progesterone-induced phosphorylation of the S308 residue, missing in neoGATA3, is involved in the proteasome-dependent degradation of GATA3, consistent with our observations [25]. Furthermore, neoGATA3 was able to enter the nucleus in the absence of the endogenous GATA3 (Figure 4b and Supplementary Figure 6d-e). The C-terminal Zn finger domain of GATA3 mediates DNA binding, while the N-terminal Zn finger stabilizes the binding and is important for the interaction with co-factors [26]. Unlike wtGATA3, purified neoGATA3 - which lacks the C-terminal Zn finger - showed only a weak binding to an oligonucleotide containing two palindromic GATAA motifs in an EMSA assay (Figure 4c). Accordingly, neoGATA3 was unable to modulate the activity of the *CDH1* and *CDH3* promoters, two known GATA3 targets [27, 28], in HEK293 cells using luciferase reporter assays (Figure 4d). Stable expression of wtGATA3 in BT20 cells resulted in reduced proliferation (P=0.054) and BrdU incorporation (P=0.012) (Figure 4e) while expression of neoGATA3 did not (Figure 4e). Similar results were obtained in MDA-MB-468 cells (Supplementary Figure 6f). These findings indicate that, in GATA3-negative breast cancer cells, neoGATA3 is unable to recapitulate the transcriptional and biological effects of wtGATA3.

**Figure 4:**
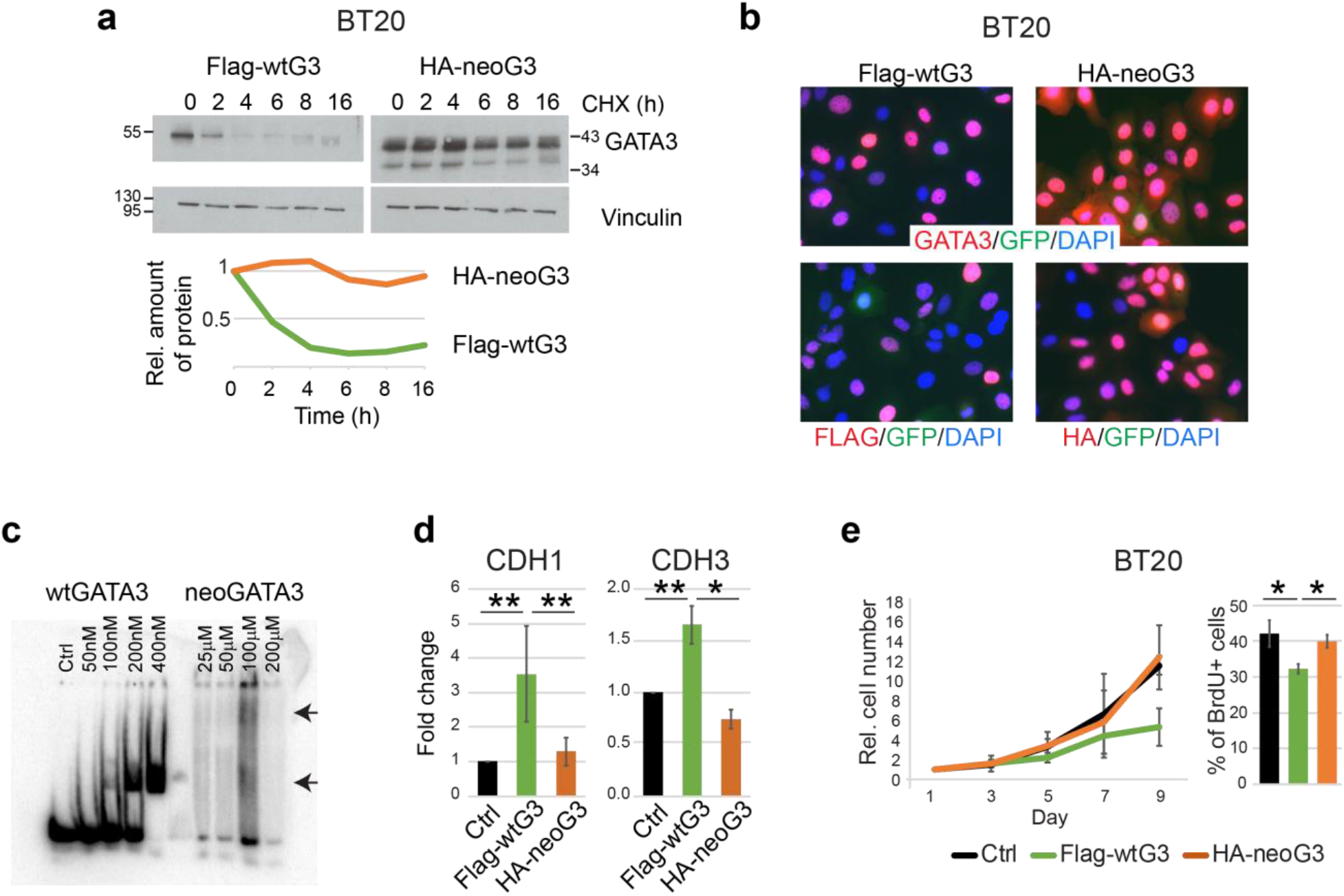
Biochemical and functional characteristics of neoGATA3 differ from wtGATA3. (**a**) Western blot showing the expression of wtGATA3 or neoGATA3 upon gene transduction in the GATA3-negative BT20 cells, after treatment with cycloheximide (CHX) for the indicated time. Vinculin was used as loading control. Quantification of relative band intensity is shown at the bottom. Images are representative of at least three independent experiments. (**b**) Immunofluorescence using the N-ter GATA3 antibody (top panels) or tag-specific antibodies (bottom panels, left: Flag, right: HA) in BT20 cells expressing either Flag-wtG3 or HA-neoG3, as indicated. DAPI was used to counterstain nuclei, GFP was expressed by the lentiviral vector used for the transduction. (**c**) EMSA assay performed with recombinant wtGATA3 or neoGATA3 and DNA fragment containing two GATAA motifs. (**d**) Luciferase reporter assay using the promoter regions of either *CDH1* or *CDH3* upstream of the luciferase cDNA. HEK293 cells were transiently transfected with the indicated constructs and luciferase activity was measured after 48h. A GFP-expressing plasmid was co-transfected to normalize for transfection efficiency by western blotting (not shown). (**e**) Growth curve (left) and percentage of BrdU+ cells (right) measured in BT20 cells transduced with the indicated constructs. Data are represented as mean ± standard deviation of at least three independent experiments. Two-sided Student’s T test *P<0.05, **P<0.01.

### NeoGATA3-expressing ER+ breast cancer cells show enhanced epithelial differentiation

Because neoGATA3 mutations are exclusively found in ER+ tumors (Supplementary Figure 3b), we assessed the function of neoGATA3 in T47D and ZR75-1, two ER+/GATA3+ BC cell lines (Figure 5a). Exogenous neoGATA3 protein was more stable also in this context and the stability of endogenous GATA3 was not affected by the mutant (Figure 5b): the half-lives of endogenous wtGATA3 and neoGATA3 were approximately 2h vs. >8h. This is consistent with the observation that, in the TCGA-BRCA series, total GATA3 protein levels were significantly higher in the neoGATA3-mutant vs. WT tumors (P=1.72e-07) [3] (Figure 5c). Overexpression of wtGATA3 or neoGATA3 had no significant effect on the proliferation and would healing capacity of T47D and ZR75-1 cells (Supplementary Figure 7a, b). Ectopic expression of neoGATA3 in ZR75-1 cells to a higher level than the endogenous protein led to a modest, significant, up-regulation of CDH1 mRNA (P=0.05) and reduced expression of VIM (P=0.036) and CDH3 (P=0.017) (Figure 5d) compared to both control- and wtGATA3-transduced cells, while other differentiation markers were not significantly changed (Supplementary Figure 7c). T47D cells, where the ectopically expressed neoGATA3 was roughly as abundant as the endogenous protein, showed a tendency to higher expression of CDH1 (Supplementary Figure 7c). This is consistent with previous work showing that GATA3 can inhibit the epithelial-to-mesenchymal transition (EMT) and the expression of basal markers in basal-like BC cells [27, 28]. Most importantly, our data are consistent with RPPA data from the TCGA-BRCA cohort, indicating a modest but significantly higher expression of CDH1 (P=0.0024) in neoGATA3-mutant tumors (Figure 5e). The CDH1 mRNA was not significantly changed in the METABRIC or TCGA-BRCA tumors carrying neoGATA3 mutations, suggesting post-transcriptional regulation of E-cadherin, while other differentiation- and EMT-related genes appeared to be significantly modulated in both cohorts (Supplementary Figure 8a-b). Altogether, these data suggest that neoGATA3 favors a well-differentiated phenotype in ER+ tumors, which might partly explain the associated good prognosis.

**Figure 5:**
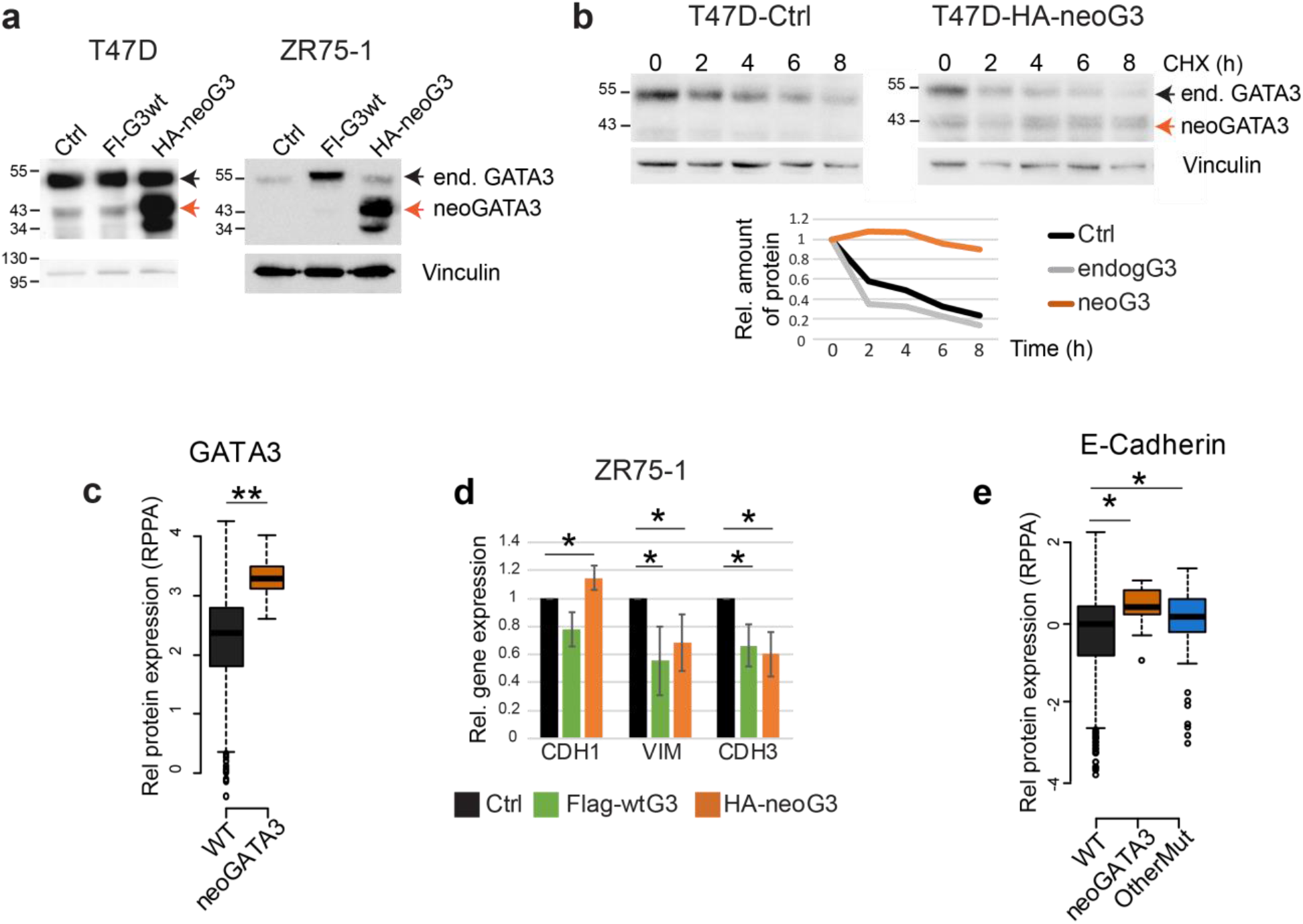
NeoGATA3 overexpression in GATA3+/ER+ cells favours an epithelial phenotype. (**a**) Western blot showing ectopic expression of wtGATA3 (Fl-G3wt) or neoGATA3 (HA-neoG3) in T47D and ZR75-1 cells. Vinculin was used as loading control. (**b**) Western blot showing the expression of endogenous GATA3 or ectopically expressed neoGATA3 in T47D cells treated with cycloheximide (CHX) for the indicated time. Vinculin was used as loading control. Quantification of relative band intensity is shown at the bottom. The endogenous GATA3 band was quantified as well in the neoGATA3-transduced cells. The images are representative of at least three independent experiments. (**c**) RPPA data from the TCGA cohort showing GATA3 expression levels in tumors of the three groups of patients (WT n=533, neoGATA3 n=18). (**d**) RT-qPCR data showing the expression of the indicated genes in ZR75-1 cells transduced with the indicated constructs, relative to Ctrl-transduced cells. Data are represented as mean ± standard deviation of at least three independent experiments. (**e**) RPPA data from the TCGA cohort showing E-cadherin expression levels in tumors of the three groups of patients (WT n=533, neoGATA3 n=18, OtherMut n=75). Two-sided Student’s T-test *P<0.05, **P<0.01.

### Genomic binding of ER is reduced in cells expressing neoGATA3

The patient-based transcriptomics analyses and the effects observed in ER+ cells suggested that neoGATA3 modulates the ER-dependent program. Treatment of hormone-depleted control T47D cells with low dose 17β-estradiol (E2) led to increased proliferative activity at 24-72h, which was blunted by 4OH-Tamoxifen (TMX). By contrast, cells overexpressing neoGATA3 showed a significantly lower response to E2 addition (P=0.025) (Figure 6a,b). A slight tendency to a reduced increase in proliferation was observed also in wtGATA3-overexpressing cells (Figure 6a-b). Similar, although less prominent, findings were made using ZR75-1 cells (Supplementary Figure 9a-b).

**Figure 6:**
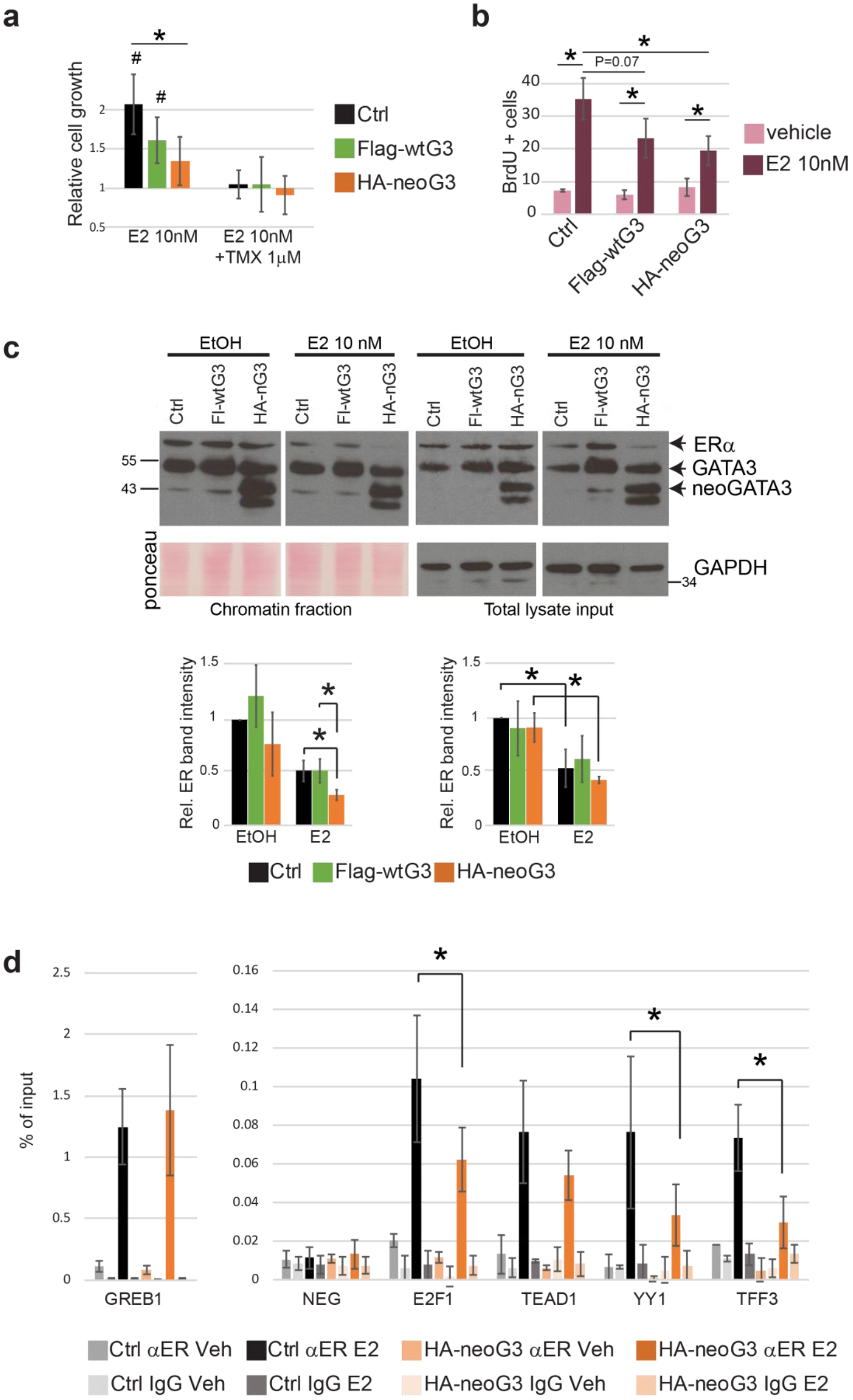
NeoGATA3 reduces the binding of ER to chromatin. (**a**) Graphs showing the relative cell growth of T47D (top) and ZR75-1 (bottom) cells transduced with the indicated constructs, after 48h in hormone-depleted (HD) medium followed by 72h of treatment with E2 (10nM) or with E2 (10nM) and TMX (1μM). All values are normalized to vehicle-treated cells of each experimental group. Data are represented as mean ± standard deviation of at least three independent experiments. *P<0.05 compared with treated Ctrl cells, #P<0.05 compared to vehicle-treated cells of the same experimental group. (**b**) Graphs showing the percentage of BrdU-positive nuclei in T47D cells transduced with the indicated constructs, after 48h in HD medium followed by 24h treatment with E2 (10nM) or with vehicle. Data are represented as mean ± standard deviation of at least three independent experiments. (**c**) Western blots showing total (right) and chromatin-bound (left) ER and GATA3 in T47D cells transduced with the indicated constructs and treated as indicated for 24h after 48h in HD medium. The Ponceau red-stained membrane was used as loading control for the chromatin fraction, GAPDH was used for the input. The images are representative of three independent experiments. The quantification of the ER band intensity is shown below. (**d**) ChIP for ER in T47D cells transduced with the indicated constructs and treated with vehicle or E2 (10nM) for 24h. Enrichment of ER on the indicated regions was measured by qPCR and represented as % of the input. NEG= gene-less region. Data are shown as mean ± standard deviation of at least three independent experiments. Two-sided Student’s T test *P<0.05.

To understand the mechanisms through which neoGATA3 interferes with the ER-dependent program, we first checked whether neoGATA3 expression affected the modulation of ER protein upon hormone starvation and subsequent stimulation with E2 or TMX, which induce a reduction and an increase of ER, respectively [17]. Expression of neoGATA3 did not affect the total ER levels in any of the tested conditions, both in T47D and ZR75-1 cells (Figure 6c and not shown). We then checked the chromatin-bound portion of the receptor after cross-linking. While this was unchanged in hormone-starved T47D cells expressing neoGATA3, its reduction after stimulation with 10nM E2 was significantly more pronounced compared to both control-transduced cells (P=0.036) and cells overexpressing wtGATA3 (P=0.016) (Figure 6c). Both endogenous GATA3 and neoGATA3 were detected at similar levels in the chromatin fraction in hormone-starved cells as well as after E2 stimulation in the three cell populations (Figure 6c).

To assess ER binding to its target genes in T47D cells, we used chromatin immunoprecipitation (ChIP) followed by qPCR. We analyzed a set of known ER binding sites based on the ENCODE/HAIB ERα ChIP-Seq experiment in T47D treated with 10nM estradiol [29] and the “ER core binding events” defined by Ross-Innes et al [24]. The *GREB1* locus shows a prominent ER peak in the ENCODE ChIP-Seq, which is among the ER core binding events. ER binding on this region was equally induced in control vs. neoGATA3-transduced cells after 24h E2 stimulation (Figure 6d). On the other hand, ER peaks of lower intensity were reported on enhancer/promoter regions close to *E2F1*, *TEAD1*, *YY1*, and *TFF3*, which were not among the ER core binding events. ER binding was induced on all these regions in control T47D cells after 24h E2 stimulation, however the induction in neoGATA3-transduced cells was significantly lower for the binding sites close to *E2F1* (P=0.012), *YY1* (P=0.006) and *TFF3* (P=0.025) and a tendency to reduced binding was observed at the region close to *TEAD1* (Figure 6d). Interestingly, DNA binding motifs for E2Fs, TEAD1, and YY1 were significantly enriched in the promoters of differentially expressed genes in the neoGATA3 METABRIC patients (Figure 3c).

Altogether, these data indicate that neoGATA3 interferes with the binding of ER to chromatin upon estrogen stimulation, especially on target regions that are weakly bound in normal conditions, contributing to reduced downstream transcriptional output.

### NeoGATA3 interferes with progesterone-induced growth arrest

While the essential role of GATA3 for ER activity is well known, its relation with PR is much less studied. NeoGATA3 appeared to interfere with the transcriptional response to progesterone in tumors, especially in pre-menopause (Figure 3f and Supplementary Figure 5d). The S308 residue, which is phosphorylated upon progesterone stimulation and signals to the proteasome [25], is absent from neoGATA3. Consistently, expression of endogenous GATA3 - but not of neoGATA3 - was reduced after treatment with 100nM progesterone for 24h (Figure 7a). In estrogenic conditions (i.e. in medium containing FBS) progesterone induced growth arrest in T47D cells, measured as BrdU incorporation after 24h of treatment and as cell viability after 6 days (Figure 7b-c). This growth arrest was significantly reduced in neoGATA3-expressing cells both at 24h (P=0.041) and after 6 days (P=0.007). Interestingly, cells overexpressing wtGATA3 showed an intermediate phenotype in the 6-day exposure, (Figure 7c) possibly due to the overexpression of the protein from an ectopic promoter, evading the PR-dependent transcriptional inhibition [25].

**Figure 7:**
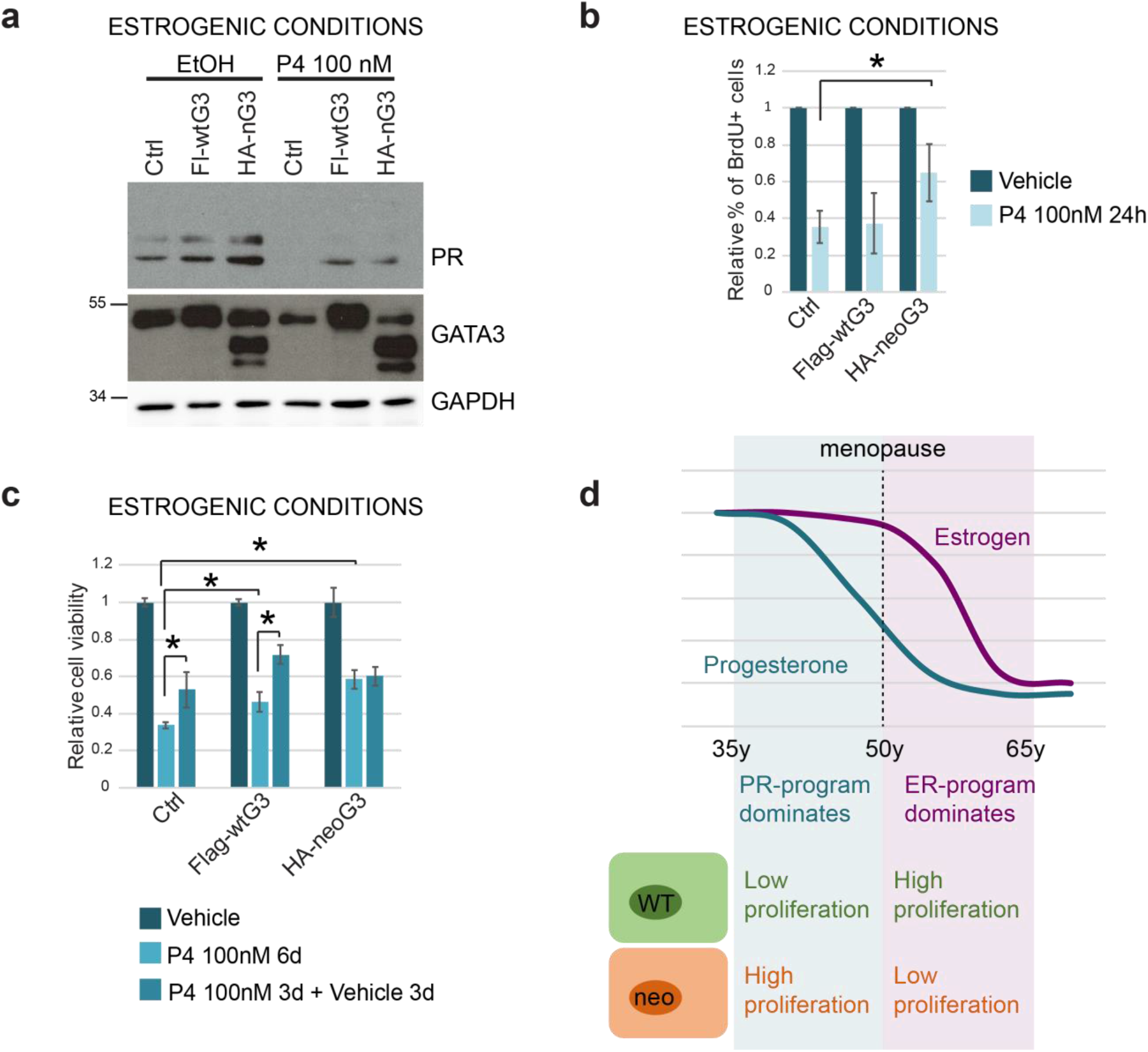
NeoGATA3 interferes with the PR-dependent growth arrest. (**a**) Western blot showing expression of PR, wtGATA3, and neoGATA3 in T47D cells transduced with the indicated constructs and treated with progesterone (P4) (100nM) for 24h in normal medium. GAPDH was used as loading control. (**b**) Graph showing the relative percentage of BrdU+ cells in the indicated cell population after 24h treatment with vehicle or 100nM P4. (**c**) Graph showing the relative cell viability measured with crystal violet staining of the indicated cell populations after vehicle-treatment, 6 days treatment with 100nM P4, or 3 days with 100nM P4 followed by additional 3 days in normal medium. Cells were kept in normal medium containing hormones. In (**b**) and (**c**) the results are normalized to the respective vehicle control and are shown as mean ± standard deviation of at least three independent experiments. Two-sided Student’s T test *P<0.05. (**d**) The working model: neoGATA3 confers cells with a proliferative advantage when both estrogen and progesterone levels are high and the anti-proliferative PR-dependent program prevails (pre-menopause). After menopause, progesterone levels drop and the ER-dependent program dominates. In this context, neoGATA3 blunts the ER program, thus inhibiting the proliferative capacity of breast cancer cells and therefore the tumor relapse. This might explain why neoGATA3 mutations are selected and why they correlate with good prognosis in patients.

In order to mimic the hormonal changes associated with pre- vs. post-menopausal status, we cultured T47D cells in an estrogen-high/progesterone-high medium (E-hi/P-hi) for 3 days and then in an estrogen-high/progesterone-low medium (E-hi/P-lo) for an additional 3 days. Both control and wtGATA3-overexpressing cells were arrested in E-hi/P-hi conditions but partially recovered proliferation after changing to E-hi/P-lo conditions. On the contrary, neoGATA3 cells remained arrested in E-hi/P-lo conditions (Figure 7c). These data support a model whereby neoGATA3 interferes with both the ER- and the PR-dependent programs, but its effects are contingent on the combined hormonal context (Figure 7d).

## Discussion

The recent large-scale sequencing efforts have brought *GATA3* to the fore as one of the most commonly mutated genes in BC [2, 3, 4]. *GATA3* was shown to be a paradigm of how genetic alterations in a given gene should not be lumped into a single class [5, 6]. Our work adds an important concept to this, namely the time/age-dependent effect of driver mutations: selected in a specific context during tumor evolution, they might have different functions later on. This notion calls for an exhaustive functional characterization to refine our understanding of their action and improve clinical application, especially when searching therapeutic targets.

The *GATA3* X308_Splice mutation, affecting the splicing between exons 4 and 5 produces a mutant protein that lacks the second ZnFn and carries a unique 44 aa peptide (neoGATA3). Importantly, we identify 5 additional mutations producing a protein with a partially or fully identical C-terminal peptide, supporting a strong selection for neoGATA3. NeoGATA3 mutations are associated with less aggressive tumors and better outcome in patients. NeoGATA3 interferes with the genomic binding of ER upon estrogen stimulation, possibly blunting its E2F-mediated proliferative function. In addition, neoGATA3 interferes with the PR-dependent anti-proliferative program in a context of high estrogen/high progesterone, mimicking pre-menopausal status. Therefore, neoGATA3 mutations represent context-specific driver mutations.

Our data provide experimental evidence to the recent prediction that the X308_Splice mutation produces a mutant transcript lacking 7nt [7] and confirms that a shorter GATA3 protein, consistent with the size of neoGATA3, which was observed in one tumor carrying the X308_Splice mutation was indeed its product [14]. A recent paper identified the mutant transcript in the RNA-Seq data from the TCGA-BRCA cohort [21], which we confirm and extend here. The neoGATA3-specific antibody recognizes the predicted mutant protein. NeoGATA3 is more stable than the wild type protein, possibly because it lacks S308 that targets wtGATA3 for degradation [25]. The lack of second GATA3 Zn finger in neoGATA3 accounts for the weaker binding to DNA in the absence of the wild type protein. Preliminary exploration of the predicted DNA binding motifs of the neoGATA3 protein with Motif Scan (not shown) suggested the possibility that the first residues of the neopeptide, together with the N-terminal Zn finger domain, generate a longer DNA binding domain, possibly with distinct specificity of binding. Further studies are warranted, to address these issues in detail. Importantly, the neoGATA3 protein often showed heterogeneous expression in tumor sections, while a GATA3 protein was always detected in 100% of cells with an N-ter directed antibody, indicating that neoGATA3 mutations are likely heterozygous and the wild type protein is co-expressed. Furthermore, we observed large amounts of neoGATA3 in the cross-linked chromatin fraction in T47D cells, suggesting that neoGATA3 might be in a complex with the wild type protein and/or other DNA-binding factors. It was suggested that GATA factors can bind adjacent motifs as dimers [30] but a direct proof that they are in the same complex in cells is missing. We have not observed wtGATA3 in the same complex with neoGATA3 in co-immunoprecipitation experiments (not shown) but we cannot exclude that they might form dimers only on the DNA.

To understand the function of neoGATA3 mutations in breast cancer, two major questions were addressed, namely: 1) what is the mechanism underlying the better prognosis linked to neoGATA3 mutations in patients; 2) why is a mutation associated with better prognosis selected in tumors.

### Association of neoGATA3 mutations with good prognosis

The unique C-terminal peptide of neoGATA3 is a predicted neoantigen [21] which was proposed to be associated with increased immune response, mostly mediated by T-cells [31]. A neoantigen-elicited tumor clearance by the immune system would be an appealing explanation for the better outcome observed in patients carrying the neoGATA3 mutations. However, we did not find evidence for this, using multiple methods of analysis. We observed that the tumor immune milieu is indeed altered in the neoGATA3 tumors, but the exact contribution of the immune infiltrates in the neoGATA3-dependent phenotype and whether/how neoGATA3 directly or indirectly influences the tumor microenvironment remains to be understood.

Another potential explanation of neoGATA3 being associated with good outcome stems from the known pro-differentiation and anti-EMT functions of wild type GATA3 [9][13, 28]. Tumors expressing neoGATA3 showed a modestly more pronounced epithelial phenotype, and overexpression of the neoGATA3 had some anti-EMT functions in ER+ cells. However, the observed differences were only minor, therefore the enforced differentiation is likely not the major driver of the neoGATA3-dependent good outcome.

ER has a well-described oncogenic function in breast cancer, which has been clinically exploited for long time. Since GATA3 is a crucial ER co-factor [11], and multiple ER-related pathways were significantly modulated in neoGATA3 mutant tumors, we explored the possibility that neoGATA3 interferes with the ER-dependent transcriptional program and showed that this is indeed the case, possibly explaining the association with outcome. GATA3 and ER regulate each other and GATA3 silencing in breast cancer cells reduces the total ER levels, abrogating the burst of proliferation induced by estrogen [10]. NeoGATA3 reduced, but did not fully abrogate, the response to estrogen. ER mRNA and protein were not markedly reduced in the cells or in patients, suggesting that neoGATA3 might exert a partial dominant-negative effect on the wild type protein, through the interference with the availability of co-factors. It was also shown that the GATA3 pioneering activity is required at a subset of inactive enhancers to open the chromatin and allow the ER to bind. When GATA3 is silenced, ER is redistributed on the chromatin upon estrogen stimulation, changing the transcriptional outcome [11]. In the presence of neoGATA3, we observed a striking reduction of the global amount of ER bound to the chromatin, although the levels of H3K27ac, a marker of open chromatin, were not significantly reduced at multiple known ER binding sites (checked by ChIP-qPCR, not shown) suggesting an impaired recruitment of ER. As the C-terminal part of GATA3 is thought to be responsible for protein-protein interactions, the novel peptide contained in neoGATA3 could quench the formation of a functional ER transcriptional complex. Future genome-wide analysis of ER binding, as well as proteomics analysis of the ER-GATA3 and ER-neoGATA3 complexes will provide further details about the precise regulation of the ER chromatin binding.

ER regulates E2F expression both in an estrogen-dependent [32, 33] and -independent manner [34]. In particular, the ligand-independent activation of an E2F transcriptional program mediated the resistance to estrogen deprivation of ER-dependent BC cells, and the expression of an E2F-dependent signature was associated with worse response to short term aromatase inhibitor treatment in patients [34]. Our data indicate that the E2F-driven program is blunted in the tumors carrying neoGATA3 mutations, possibly through the lower ER binding to chromatin. Since most patients treated with endocrine therapy initially respond but eventually develop resistance and succumb to the disease, our observations suggest that tumors expressing neoGATA3 are less prone to develop resistance, consistent with a longer patient survival.

### Context-specific driving activity of neoGATA3

Having unraveled one mechanism through which neoGATA3 confers a less aggressive behavior to breast cancer cells, we were puzzled by the fact that the X308_Splice mutation is highly selected during tumor evolution. We reasoned that there might be a context in which neoGATA3 gives a proliferative advantage to the cells. NeoGATA3 mutations are highly frequent among pre-menopausal METABRIC patients (21/253, 8.9%), in whom the levels of estrogens and progestogens are relatively high and ER and PR have antagonistic effects through the modulation of shared targets (genomic agonism) [35]. GATA3 and PR are in the same protein complex [36]. However, the functional relation between both proteins and the link to ER are still to be investigated. Furthermore, PR dampens the activity of GATA3 both at transcriptional and post-translational levels [25]. The strong selection of a mutation abrogating this residue points to a function in evading the PR-dependent anti-proliferative program as we observed in T47D, where progesterone-induced growth arrest was less prominent in neoGATA3-expressing cells. Interestingly, wtGATA3 expression in T47D cells from an ectopic promoter evading PR-dependent transcriptional inhibition only showed a tendency to reduce progesterone-induced growth arrest and 4 of the 5 neoGATA3-like mutations that we describe here retain the S308 residue and are therefore likely degraded in response to progesterone as the wild type. This would indicate that the increased stability of neoGATA3 is not the only explanation for the interference with PR function. Our data suggest that GATA3 is a crucial co-factor for both ER and PR and further –omics studies should be performed to assess its precise role in the transcriptional programs controlled by them.

Intriguingly, the wtGATA3 was expressed at much lower levels than the neoGATA3 in our experiments in T47D cells, yet wtGATA3-transduced cells showed some intermediate phenotypes between Ctrl-transduced and neoGATA3-expressing cells. This would suggest that an increase in GATA3 expression is sufficient to disrupt the balance between ER and PR programs, and that neoGATA3 is less efficient than the wild type protein at doing this. In addition, the C-terminal neopeptide of neoGATA3 may interfere - but not abolish - the formation of functional ER and PR complexes. Therefore, neoGATA3 behaves as a weak, inefficient, oncogenic driver.

In conclusion, our data suggest that neoGATA3 mutations are specifically selected in a molecular context whereby estrogen-driven mitogenic phenotypes are counterbalanced by progesterone-driven antiproliferative effects. In this particular scenario, the net output of neoGATA3 interference with both ER and PR programs is a proliferative advantage. After menopause, progesterone levels drop rapidly, so that the ER-dependent program is not compensated by the PR program and becomes the dominant pathway. In this context, neoGATA3 confers a proliferative disadvantage to the cells and is associated with better patient outcome (Figure 7d). The neoGATA3 mutations therefore result in a partial LOF and represent a subtype of context-dependent driver mutations associated with distinct clinical features.

## Acknowledgments

We want to thank G. Timelthaler, M.A. Quintela and F. Reyal for valuable contributions, J. Valcárcel, J. Muñoz-Pérez, and O. Domínguez for discussions.

## Methods

### Patient-related information

Data from the TCGA-BRCA cohort (clinical, gene expression, RPPA, and mutations) were obtained from the UCSC XENA browser (www.xena.ucsc.edu), while the data from the METABRIC cohort (clinical, gene expression, and mutations) were obtained from the cBio Cancer Genomics Portal (www.cbioportal.org). Survival analyses were performed with R; LogRank statistics was calculated with the Cox proportional hazard method. The MCP counter values were obtained from a previously published work [22]. The number of samples included in each analysis varies because not all samples had full clinical data and expression data.

### Patient samples

Formalin-fixed paraffin-embedded sections of resected breast tumors, as well as DNA, RNA, and protein lysates from fresh tissue, together with the corresponding clinical data, were obtained from a cohort of 102 patients receiving surgery in the Hospital Clínico Universitario de Valencia/INCLIVA and one additional cohort of 100 patients from the Hospital Val d’Hebron (Barcelona). FFPE from TMAs and full sections, as well as genomic DNA from one patient were obtained from the METABRIC cohort as described previously. All procedures were approved by the institutional Ethics Committees and informed consent was obtained from all patients.

The presence of the X308_Splice DNA mutation was assessed by PCR of the intron4-exon5 junction, followed by Sanger sequencing.

### Cell lines and treatments

All cell lines used here are commercially available from ATCC. Cells were cultured in standard conditions (37 °C, 20% O_2_, 5% CO_2_) and periodically checked for mycoplasma contamination through PCR. HEK293FT cells were maintained in High Glucose DMEM supplemented with 10% FBS and 1% antibiotics (Pen/Strep). BT20, MDA-MB-468, T47D, and ZR75-1 were maintained in RPMI supplemented with 10% FBS and 1% antibiotics (all from Sigma-Aldrich). For the treatment with E2 and TMX in hormone-depleted medium, cells were kept for at least 48h in RPMI without phenol red, supplemented with 10% of charcoal-stripped FBS. T47D and ZR75-1 were authenticated by Eurofins Genomics. 17β-estradiol (E2), 4OH-Tamoxifen (TMX), progesterone (P4), cycloheximide (CHX), and the proteasome inhibitor MG132 were purchased from Sigma-Aldrich and dissolved in EtOH, which was then used as vehicle control. 5-Bromo-2’-deoxyuridine (BrdU) was purchased from Sigma-Aldrich, dissolved in water, and added to the cells for 2-4 hours at 50µM.

### Plasmids, transfection and lentiviral transduction

The wild type GATA3 cDNA was a kind gift of C.M. Perou and J. Usary, the neoGATA3 cDNA was generated from it through site-directed mutagenesis. The CDH1 and CDH3 promoter reporter plasmids were a kind gift of A. Muñoz and J. Paredes, respectively. Promoter reporter plasmids were transfected into HEK293FT cells using jetPRIME DNA transfection reagent (Polyplus) following the instructions of the manufacturer. The pEGFP-C1 plasmid (Invitrogen) was co-transfected at 1:10 ratio for normalization. GATA3- and neoGATA3-expressing lentiviral plasmids were transfected into HEK293FT cells together with packaging plasmids with calcium-phosphate precipitation. Virus-containing supernatant from HEK293FT packaging cells was collected, filtered (0.45µm) and used to transduce epithelial cells.

### RNA-Seq re-analysis and identification of the neoGATA3 transcript

Raw RNA-seq data were downloaded from CGHUB. RNA sequencing consisted of 48-50bp paired-end reads. To align reads to the human genome (GRCh37/hg19) TopHat-2.0.10 was used with Bowtie 1.0.0 and Samtools 0.1.19, allowing two mismatches and five multihits. Transcript assembly, estimation of abundance, and merging were performed with Cufflinks 2.2.1.

### Immunoblotting

Cells were lysed with Laemmli buffer and proteins were separated by SDS-PAGE. The following primary antibodies were used: anti-GATA3 (Cell Signaling #5852, or Cell Marque L50-823), anti-ER (Santa Cruz sc-543X or sc-8002), anti-Flag (Sigma-Aldrich F1804), anti-HA (BioLegend HA.11), anti-GAPDH (e-bioscience clone clone FF26A), and anti-vinculin (Sigma-Aldrich V9264). Anti-rabbit IgG or HRP-conjugated mouse IgG (Life Technologies) were used as a secondary antibodies. The chemiluminescent signals were detected with BioRad Chemidoc system or with Amersham films. Band intensity was quantified with ImageJ or with the BioRad Image Lab software.

### Immunofluorescence

Cells were seeded on glass coverslips and fixed with 4% paraformaldehyde. After permeabilization with Triton 0.1%, coverslips were incubated with primary antibodies (GATA3, Flag, or HA), followed by Alexa-Fuor-conjugated fluorescent secondary antibodies (Life Technologies). Nuclei were counterstained with DAPI and coverslips were mounted with Agilent Fluorescent Mounting Medium. Images were acquired on a Zeiss LSM700 confocal microscope. For BrdU detection, the primary antibody (Sigma-Aldrich BU-33) was incubated together with 1µg/ml DNAse I (Sigma-Aldrich) in the presence of 6mM MgCl_2_.

### Immunohistochemistry

Formalin-fixed paraffin-embedded sections were stained following standard procedures. Briefly, sections were de-paraffinized, re-hydrated, boiled in citrate buffer pH 6.0 for antigen retrieval, and incubated with 10% H_2_O_2_ in MetOH to quench the endogenous peroxidases. Afterwards, sections were incubated with primary antibodies recognizing GATA3 (Cell Signaling), neoGATA3 (home-made), and CD8α (DAKO C8/144B) overnight at 4 °C. HRP-conjugated secondary antibodies were from DAKO. 3,3-diaminobenzidine tetrahydrochloride plus (DAB+) was used as chromogen and nuclei were counterstained with hematoxylin.

### Cell proliferation, wound healing, viability assay

To monitor cell growth, 4×10^4^ cells/well were seeded in 6-well plates in duplicates and counted every other day. For BrdU incorporation, cells were seeded on glass coverslips, treated as indicated, then fixed with paraformaldehyde and stained with anti-BrdU. For would healing experiments, cells were seeded in 12-well or 24-well plates and allowed to reach confluence. Then a scratch was done with a sterile 10µl pipette tip, the well was washed and the medium was replaced with serum-free RPMI. Microphotographs of at least 3 distinct regions of the wound were taken at the indicated time points and the open area was quantified with ImageJ. To assess cell viability upon E2 and TMX treatment, 5000 cells/well were seeded in 96-well plates in quadruplicates; the following day, the medium was changed to hormone-depleted RPMI and treatment with vehicle, E2, TMX, or the combination started after 48h. After 72h, cells were fixed with ice-cold methanol and stained with crystal violet. After extensive washing and drying, the dye was extracted with 1% SDS and the absorbance at 595 nm was measured on a Tecan microplate reader. For the treatments with P4 in estrogenic conditions, 50×10^4^ cells were seeded in 12-well plates in complete RPMI. Starting from the next day, the medium was changed daily, with the addition of either 100nM P4 or the corresponding amount of EtOH. After 72h, one set of wells was changed back to normal medium. Cells were fixed with ice-cold methanol after 6 days and stained with crystal violet. Cell viability was quantified as described above.

### Luciferase assay

HEK293T cells were transfected with *CDH1* or *CDH3* promoter reporter plasmids, together with a GFP-expressing plasmid (pEGFP-C1). At the same time, empty pCDNA3 (Invitrogen), or pCDNA3 containing either wild-type GATA3 or neoGATA3 cDNA were introduced. Luciferase activity was measured with a luminometer, using a commercial luciferin solution (Promega) as a substrate. Values were normalized for transfection efficiency by checking GFP levels using western blotting.

### Gene expression analysis

Total RNA was extracted from cells using peqGOLD TriFast (VWR) according to manufacturer’s instructions, and converted to cDNA using Revert Aid reverse transcription reagents including random hexamers (ThermoFisher Scientific). Quantitative PCR was performed using SYBR-green mastermix (Promega) and run in a Prism 7900 HT instrument (Applied Biosystems). Primers were designed using Primer3Plus and reactions were done in triplicate. All quantifications were normalized to endogenous HPRT, using the standard ΔΔCt method. Primers sequences are included in Supplementary Table 4.

### Cloning of wtGATA3 and neoGATA3 in pOPIN-B

DNA sequences encoding the wild-type GATA3 region from residue 260 to 370 (wtGATA3) and the neoGATA3 region from residue 260 till the C-terminal end (neoGATA3) were PCR amplified from the full-length genes using the following oligonucleotides:

wtGATA3_Fw: aagttctgtttcagggcccgGGCAGGGAGTGTGTGAAC
wtGATA3_Rv: atggtctagaaagctttaTCAGCTAGACATTTTTCG
neoGATA3_Fw: aagttctgtttcagggcccgGGCAGGGAGTGTGTGAAC
neoGATA3_Rv: atggtctagaaagctttaTCAGGGGTCTGTTAATATTG

Amplicons were gel-purified and ligated into a pOPIN-B vector digested with HindIII and KpnI using In-Fusion technology (Clontech). The primer sequences in low case correspond to adaptors pairing with the linearized vector. This vector adds a poly-histidine tag and a cleavage site for the protease PreScission to the N-terminus of the protein. The correct insertions of the sequences in the plasmid were verified by sequencing.

### Protein production and purification

*E coli* Rosetta (DE3) pLysS cells (Novagen) transformed with pOPIN-B plasmids encoding wtGATA3 and neoGATA3 were grown in autoinduction media supplemented with 50 µg/ml kanamycin and 34 µg/ml chloramphenicol for 8 h at 37 °C followed by 21 h at 20 °C. The cells, resuspended in buffer A (20 mM Tris-HCl pH 8, 0.5 M NaCl, 10 mM imidazole, 5% glycerol and 5 mM β-mercaptoethanol) with 0.4 mM Pefabloc (Merck), were disrupted by sonication. The clarified supernatant was applied onto a 5 ml HisTrap HP column (GE Healthcare) equilibrated in buffer A connected to an FPLC-Prime (GE Healthcare). Following extensive washing with buffer A supplemented with 34 mM imidazole, the protein was eluted with buffer A with 250 mM imidazole. The sample was dialyzed overnight in buffer B (10 mM HEPES pH 7.6, 0.1 M NaCl, 5 µM ZnSO4 and 1 mM DTT) and GST-tagged PreScission protease was added in the dialysis bag (1/20^th^ of the protein weight) to cleave the N-terminal tag. Then, the sample was loaded onto a 5 ml HiTrap-SP HP column (GE-Healthcare) equilibrated in buffer B, coupled to a 5 mL GST-Trap HP (GE-Healthcare) column to retain the protease. The untagged GATA protein eluted in a salt gradient at approximately 0.15 M NaCl and was concentrated in an Amicon 3K ultracentrifugation device (Millipore) and loaded onto a Superdex 75 10/300 column (GE-Healthcare) equilibrated in buffer B and connected to an AKTA FPLC system (GE-Healthcare). The protein eluted in a single peak and was concentrated up to 10 mg·ml^-1^ as mentioned above. The protein was supplemented with 20% glycerol, frozen in liquid nitrogen and store at −80°C. All purification steps were carried out at 4°C and sample purity was assessed by SDS-PAGE.

### Electrophoretic Mobility Shift Assay (EMSA)

The following HPLC purified DNA oligonucleotides were purchased from Metabion:

DNA1_Fw: TGTCCATCTGATAAGAC
DNA1_Rv: GTCTTATCAGATGGACA

The oligonucleotides were resuspended in TE buffer (10 mM Tris-HCl pH 8.0, 1 mM EDTA) to a final concentration of 100 µM and were radiolabeled with T4 Polynucleotide kinase (PNK, Invitrogen) in a reaction containing 1 mM ATP-γ-^32^P (Perkin Elmer) for 2h at 37°C. The reaction was diluted with TE buffer and loaded into an illustra MicroSpin G-25 Column (GE) to remove excess of free-radioactive ATP. Complementary strands were annealed by heating at 95°C for 5 min, followed by slow cooling at room temperature. ^32^P-radiolabeled double stranded DNAs were stored at −20°C. Samples for EMSA were prepared in a final volume of 20 µL, by mixing 100 nM of double stranded DNA1 or DNA2 with wtGATA3 or neoGATA3 at concentrations ranging from 0–0.4 µM, in a buffer containing 5 mM HEPES pH 7.6, 0.5 mM EDTA, 4 mM magnesium oxaloacetate, 50 mM potassium chloride, 10% glycerol, 1 mM DTT, 0.1 mg·ml^-1^ BSA and 2 µg·ml^-1^ salmon DNA. After 1 h incubation at room temperature, the samples were applied to a 6% acrylamide-TBE electrophoresis gel, ran in TBE 0.5x buffer at 150 V for 30 min at 4°C. The gel was dried and exposed to a PhosphorScreen (GE-Healthcare) that was read in a Typhoon FLA 7000 (GE). Gel analysis was performed with ImageQuant software.

### Chromatin isolation

Cells grown at 70-80% confluency were cross-linked in 1% formaldehyde at room temperature for 15 min, followed by 5 min quenching with glycine. Cells were harvested in PBS, centrifuged, and nuclei were isolated through the lysis in 0.34M sucrose/10% glycerol/10% Triton buffer followed by centrifugation. Nuclei were washed and further separated in nucleoplasm and chromatin fractions with 3mM EDTA/0.2mM EGTA buffer. The chromatin fraction recovered after centrifugation was sonicated and all fractions were resuspended in Laemmli buffer to be run on SDS-PAGE.

### ChIP

Cells were grown until 70-80% confluence, treated as appropriate, and cross-linked in 1% formaldehyde at room temperature for 10 min. After quenching, cells were harvested in PBS, nuclei were enriched with hypotonic solution, and lysed with 0.5% SDS. Chromatin was sonicated for 15 min in a Bioruptor water bath (Diagenode) using high intensity and 30”on/30”off cycles and then diluted to adjust SDS concentration to 0.1%. Immunoprecipitation was performed overnight at 4 °C with protein A/G agarose beads linked to anti-ERα (sc-543X) or rabbit IgG. Beads were washed 6x after incubation and the complex was eluted and de-crosslinked by overnight incubation at 65 °C. Eluted DNA was treated with proteinase K and purified with phenol-chloroform followed by isopropanol precipitation. The isolated DNA was then used for qPCR. Primers sequences are listed in Supplementary Table 4.

### Gene Set Enrichment Analysis

Differential gene expression was computed on the METABRIC and TCGA-BRCA datasets using the Comparative Marker Selection module of Genepattern, by comparing tumors carrying a neoGATA3 mutation and all other tumors. GSEA was then calculated on the ranked list of makers using the GSEA Preranked module and interrogating the Hallmarks and the C3-TFT Transcription Factor Targets gene set collections of the MSigDb of the Broad Institute. FDR <0.05 was considered significant.

### Neogata3-specific antibody production

A peptide was synthesized based on the predicted neoGATA3-specific C-terminal sequence (PGEQGRPVRTVRPPQPHSGGGMPMGTLSAMPVGSTTSFTILTDP), including residues 309-352, plus a final Cys. The peptide was conjugated with KLH and used to immunize rabbits. Polyclonal antibodies were obtained following standard procedures.

### Statistical analyses

The specific statistical test used for each analysis is indicated in the respective figure legend or table description. Statistical analyses were performed with R studio.

## Supplementary Material

**Supplementary Table 1:** *GATA3* mutations leading to a neoGATA3 protein (fully or partially concordant with the original X308_Splice mutant).

| chromosomal location | gene location | change | C-ter | cBioportal nomenclature |
| --- | --- | --- | --- | --- |
| chr10:8111432-8111434 | intr4-exon5 | TCA-->T | neoGATA3 (44 aa) | X308_Splice |
| chr10:8111513-8111513 | exon5 | T-->TGG | 20 aa | D335Gfs*21 |
| chr10:8106083-8106084 | exon4 | TA-->T | neoGATA3 | K302Sfs*53 |
| chr10:8111452-8111453 | exon5 | CA-->C | 40 aa | T315Rfs*40 |
| chr10:8111499-8111499 | exon5 | GA-->G | 25 aa | R329Gfs*26 |
| chr10:8111487-8111487 | exon5 | A-->AAA | 34 aa | not present (Q321Qfs*34) |

**Supplementary Table 2.** Cox model for disease-specific survival among BC patients with ER+ tumors (n=1508).

|  | Univariate analysis |  |  |  | Multivariable analysis |  |  |  |
| --- | --- | --- | --- | --- | --- | --- | --- | --- |
| Variables | HR | Lower 95% | Upper 95% | P-value | HR | Lower 95% | Upper 95% | P-value |
| Age (cont. variable) | 1.02 | 1.01 | 1.03 | <0.001 | 1.01 | 1.00 | 1.02 | 0.011 |
| Tumor size (cont. variable) | 1.02 | 1.02 | 1.02 | <0.001 |  |  |  | <0.001 |
| Stage |  |  |  |  |  |  |  |  |
| T0 | 1.00 | - | - | - | 1.00 | - | - | - |
| T1 | 0.48 | 0.35 | 0.64 | <0.001 | 0.58 | 0.43 | 0.81 | 0.001 |
| T2 | 0.84 | 0.66 | 1.07 | 0.166 | 0.72 | 0.56 | 0.94 | 0.015 |
| T3 | 1.86 | 1.23 | 2.81 | 0.003 | 1.19 | 0.72 | 1.97 | 0.485 |
| T4 | 6.25 | 3.17 | 12.30 | <0.001 | 4.75 | 2.27 | 9.92 | <0.001 |
| Grade |  |  |  |  |  |  |  |  |
| I | 1.00 | - | - | - | 1.00 | - | - | - |
| II | 1.76 | 1.16 | 2.66 | <0.001 | 1.18 | 0.73 | 1.90 | 0.501 |
| III | 2.94 | 1.94 | 4.44 | <0.001 | 1.59 | 0.98 | 2.58 | 0.058 |
| PR status |  |  |  |  |  |  |  |  |
| Neg | 1.00 | - | - | - | 1.00 | - | - | - |
| Pos | 0.66 | 0.54 | 0.79 | <0.001 | 0.92 | 0.73 | 1.16 | 0.49 |
| GATA3 status |  |  |  |  |  |  |  |  |
| WT | 1.00 | - | - | - | 1.00 | - | - | - |
| Other MUT | 0.64 | 0.45 | 0.91 | 0.014 | 0.65 | 0.43 | 0.98 | 0.040 |
| neoGATA3 | 0.26 | 0.13 | 0.52 | <0.001 | 0.46 | 0.23 | 0.94 | 0.034 |
| PAM50 subtype |  |  |  |  |  |  |  |  |
| Luminal A | 1.00 | - | - | - | 1.00 | - | - | - |
| Luminal B | 2.24 | 1.80 | 2.80 | <0.001 | 1.65 | 1.28 | 2.15 | <0.001 |
| Basal-like | 2.67 | 1.75 | 4.09 | <0.001 | 2.51 | 1.44 | 4.37 | 0.001 |
| HER2E | 2.39 | 1.73 | 3.30 | <0.001 | 1.86 | 1.20 | 2.87 | 0.005 |
| Normal-like | 1.34 | 0.95 | 1.90 | 0.096 | 1.50 | 1.00 | 2.26 | 0.048 |

**Supplementary Table 3.** Cox model for overall survival among BC patients with ER+ tumors (n=1508).

|  | Univariate analysis |  |  |  | Multivariable analysis |  |  |  |
| --- | --- | --- | --- | --- | --- | --- | --- | --- |
| Variables | HR | Lower 95% | Upper 95% | P-value | HR | Lower 95% | Upper 95% | P-value |
| Age (cont. variable) | 1.05 | 1.04 | 1.06 | <0.001 | 1.05 | 1.04 | 1.05 | <0.001 |
| Tumor size (cont. variable) | 1.02 | 1.01 | 1.02 | <0.001 | 1.01 | 1.00 | 1.02 | <0.001 |
| Stage |  |  |  |  |  |  |  |  |
| T0 | 1.00 | - | - | - | 1.00 | - | - | - |
| T1 | 0.44 | 0.36 | 0.55 | <0.001 | 0.61 | 0.48 | 0.77 | <0.001 |
| T2 | 0.81 | 0.68 | 0.96 | 0.017 | 0.78 | 0.64 | 0.94 | 0.009 |
| T3 | 1.42 | 1.01 | 2.00 | 0.046 | 1.09 | 0.72 | 1.65 | 0.686 |
| T4 | 3.95 | 2.03 | 7.70 | <0.001 | 2.54 | 1.24 | 5.19 | 0.010 |
| Grade |  |  |  |  |  |  |  |  |
| I | 1.00 | - | - | - | 1.00 | - | - | - |
| II | 1.27 | 0.98 | 1.63 | 0.011 | 1.01 | 0.74 | 1.37 | 0.96 |
| III | 1.70 | 1.32 | 2.19 | <0.001 | 1.17 | 0.85 | 1.62 | 0.32 |
| PR status |  |  |  |  |  |  |  |  |
| Neg | 1.00 | - | - | - | 1.00 | - | - | - |
| Pos | 0.76 | 0.66 | 0.87 | <0.001 | 0.88 | 0.73 | 1.04 | 0.139 |
| GATA3 status |  |  |  |  |  |  |  |  |
| WT | 1.00 | - | - | - | 1.00 | - | - | - |
| Other MUT | 0.74 | 0.58 | 0.93 | <0.001 | 0.77 | 0.58 | 1.04 | 0.089 |
| neoGATA3 | 0.33 | 0.22 | 0.51 | <0.001 | 0.58 | 0.36 | 0.92 | 0.020 |
| PAM50 subtype |  |  |  |  |  |  |  |  |
| Luminal A | 1.00 | - | - | - | 1.00 | - | - | - |
| Luminal B | 1.57 | 1.35 | 1.83 | <0.001 | 1.15 | 0.95 | 1.39 | 0.147 |
| Basal-like | 1.48 | 1.04 | 2.11 | 0.03 | 1.37 | 0.84 | 2.23 | 0.203 |
| HER2E | 1.55 | 1.22 | 1.97 | <0.001 | 1.25 | 0.90 | 1.76 | 0.183 |
| Normal-like | 0.98 | 0.76 | 1.27 | 0.90 | 1.22 | 0.89 | 1.68 | 0.206 |

**Supplementary Table 4.**
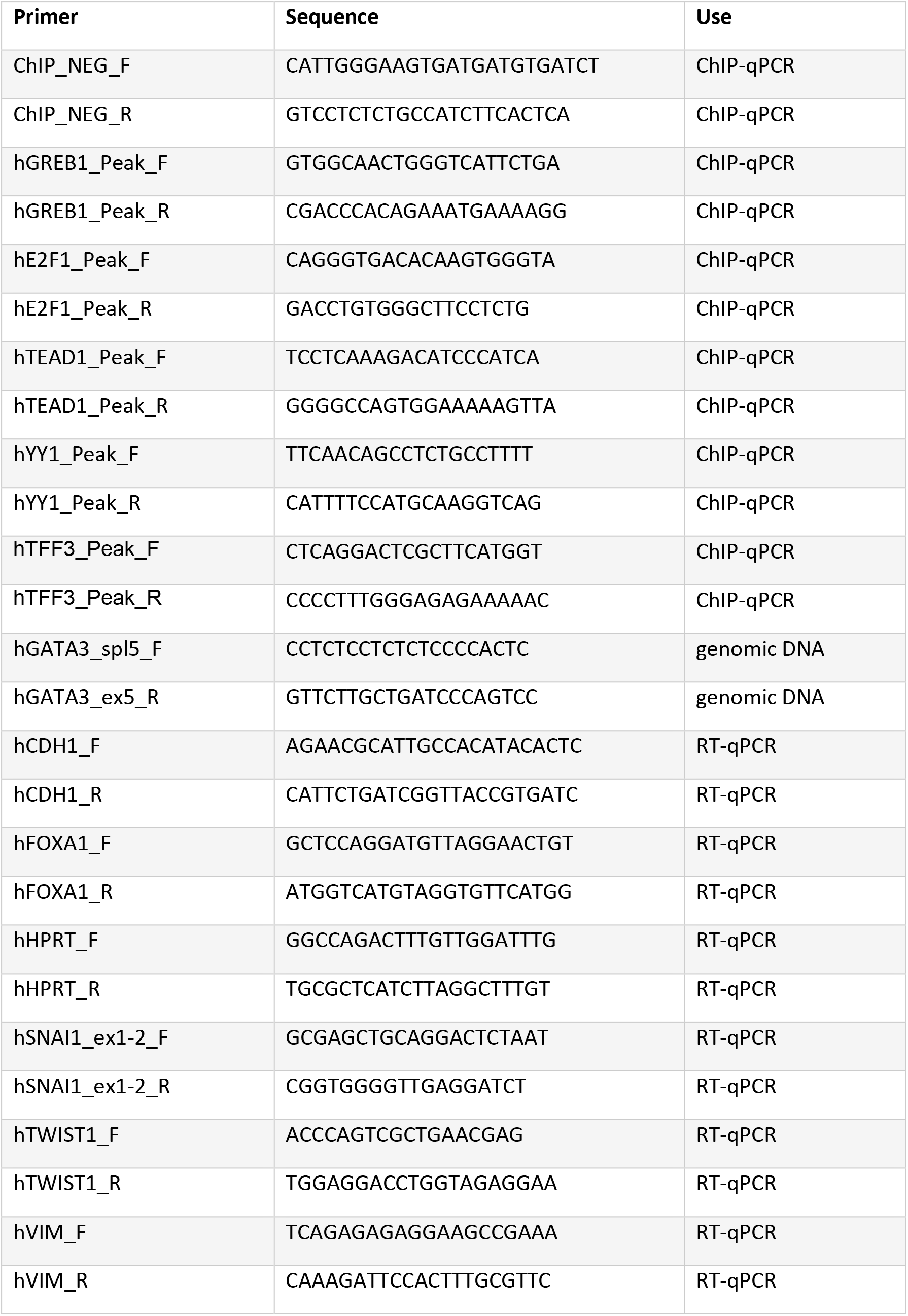
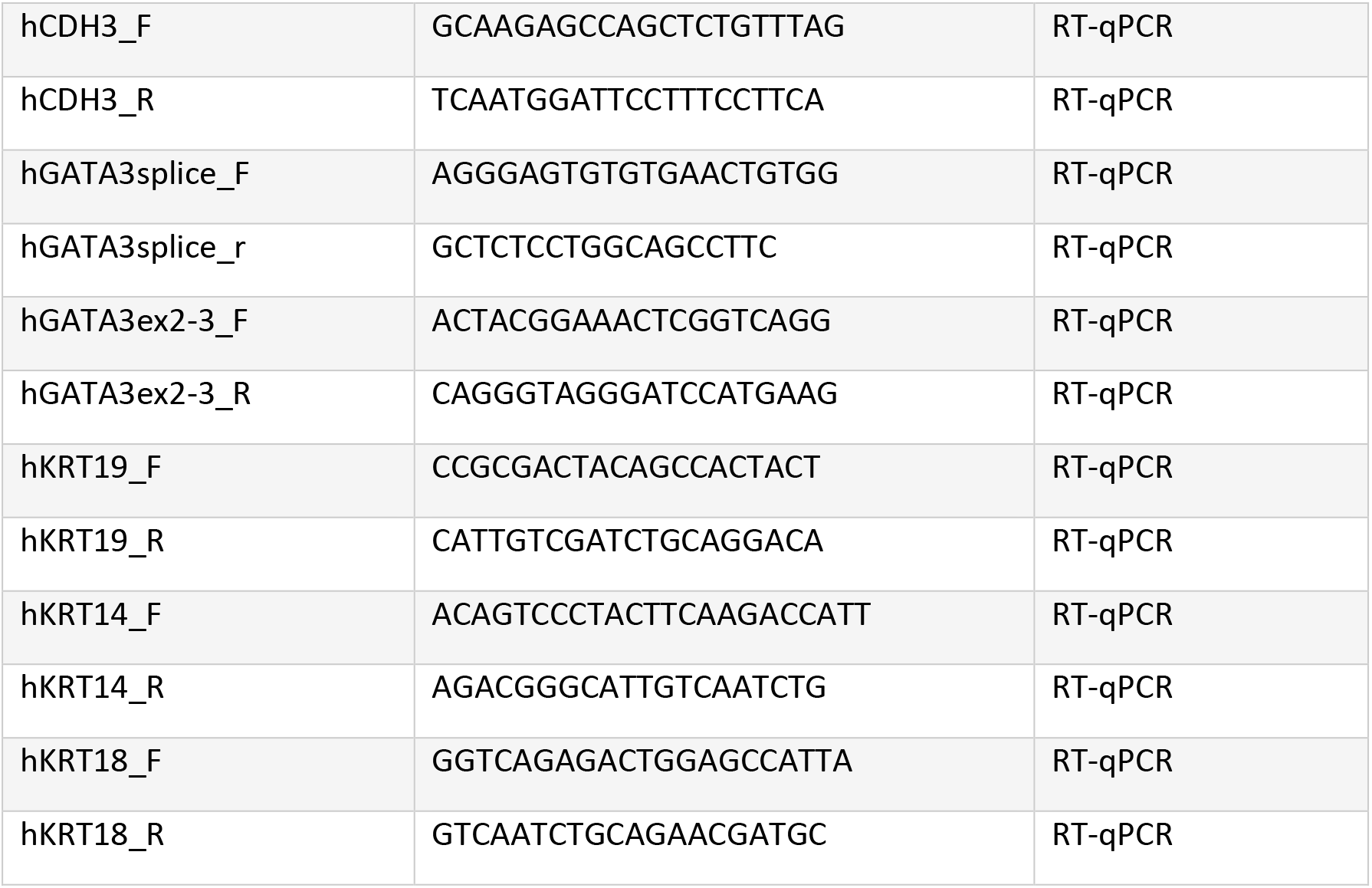
Primers used for PCR, RT-qPCR, and ChIP-qPCR.

**Supplementary Figure 1:**
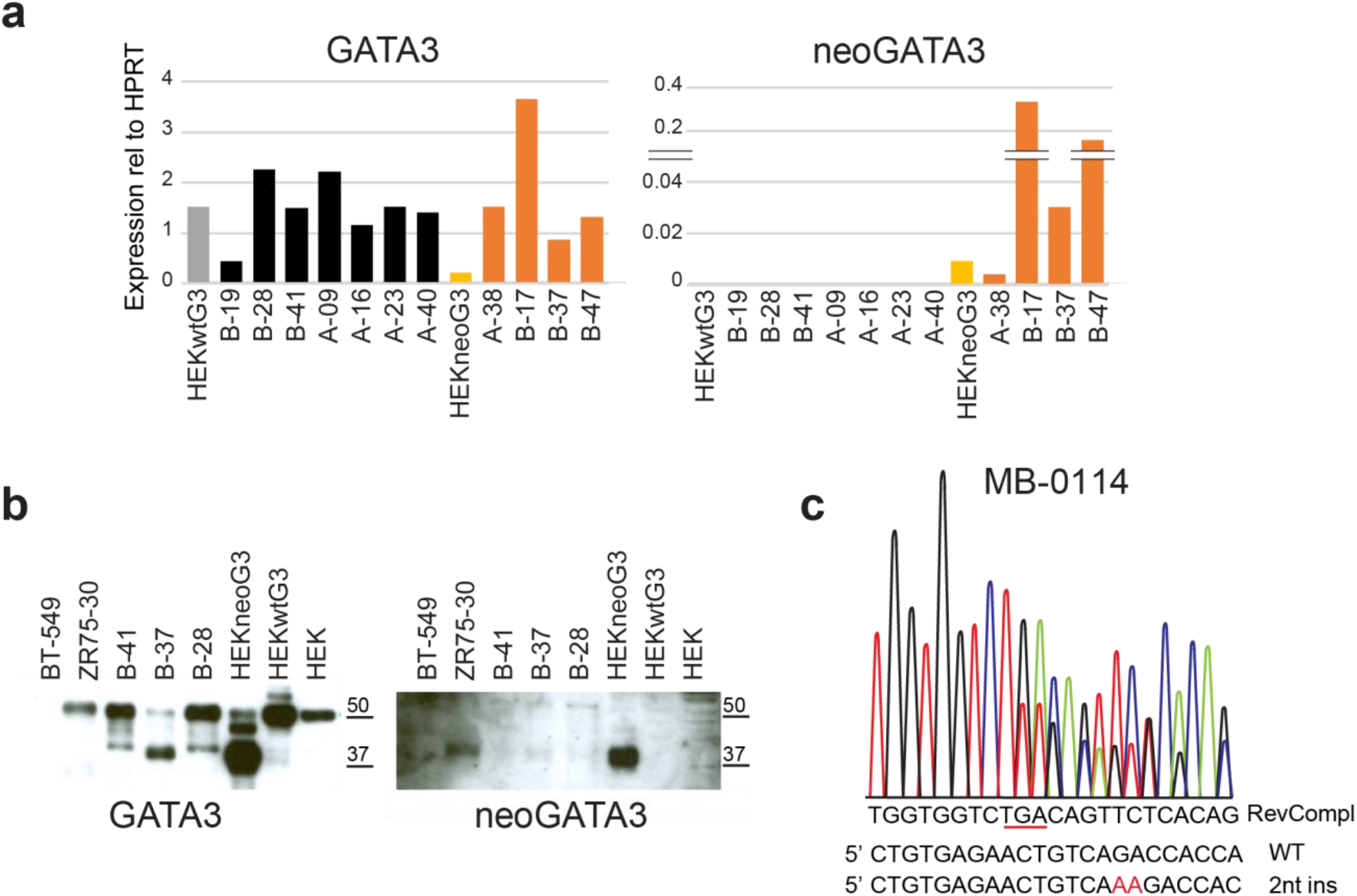
The BC-specific X308_Splice hotspot *GATA3* splice mutation produces a truncated transcript and protein. (**a**) qPCR analysis using primers amplifying both the wild type and the mutant GATA3 (left) or specific for the truncated neoGATA3 transcript (right) on tumor tissue. HEK293 cells transfected with either wtGATA3 or neoGATA3 were used as controls. (**b**) Western blot on total protein extract of tumor tissue, using an antibody that recognizes both wild type and mutant GATA3 (left) or with the neoGATA3-specific antibody (right). HEK293 cells transfected with either wtGATA3 or neoGATA3 were used as controls. (**c**) The 2-nucleotide insertion identified in the MB-0114 METABRIC tumor upon re-sequencing.

**Supplementary Figure 2:**
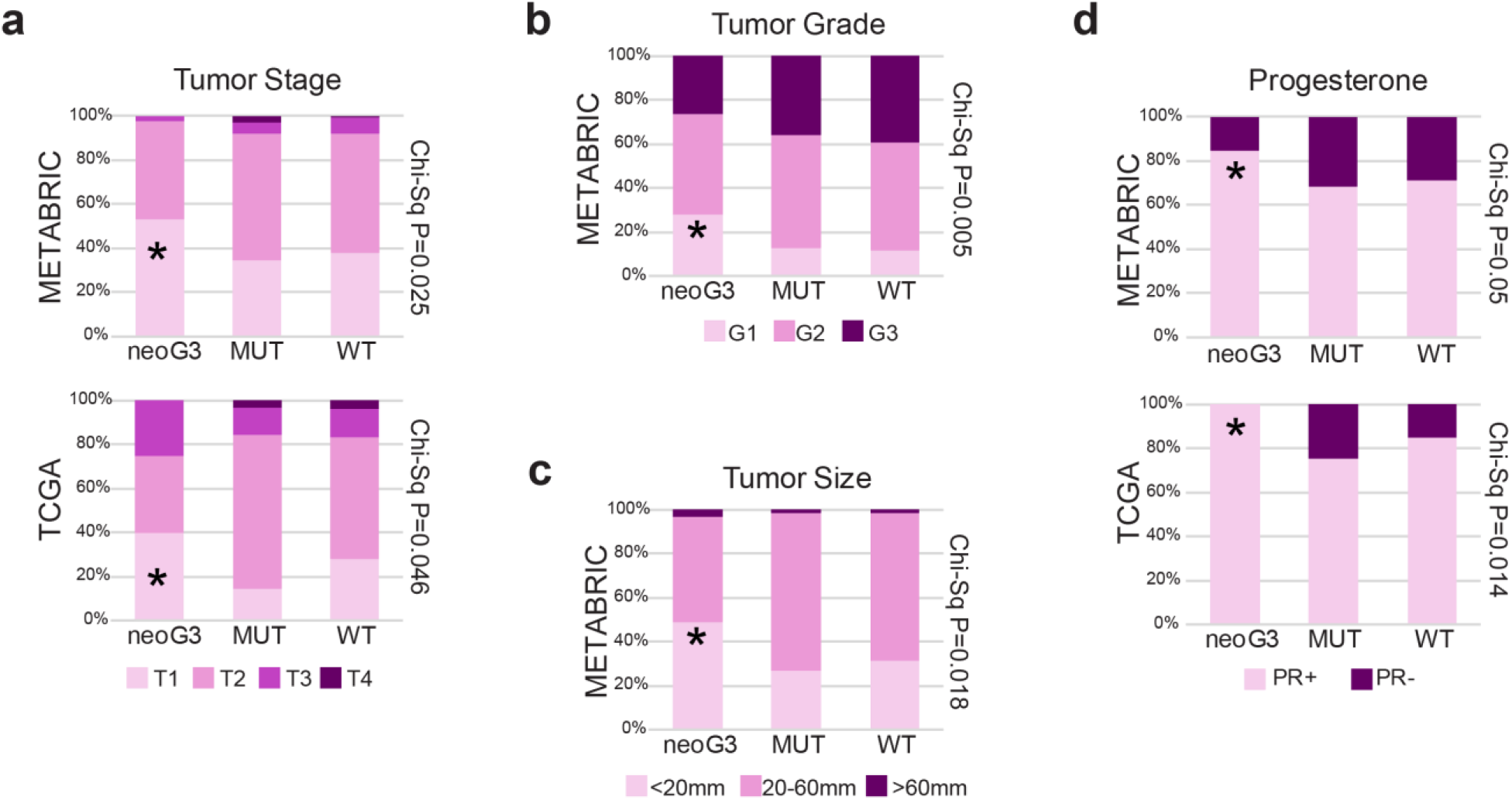
NeoGATA3 mutations are associated with markers of good prognosis. (**a-d**) Distribution of tumor stage (**a**), grade (**b**), size (**c**), and PR status (**d**) among the three groups of patients in the METABRIC and TCGA-BRCA cohorts (METABRIC: stage WT n=648, neoGATA3 n=49, OtherMut n=104; grade WT n=891, neoGATA3 n=57, OtherMut n=133; size WT n=924, neoGATA3 n=59, OtherMut n=139; PR WT n=929, neoGATA3 n=59, OtherMut n=139; TCGA-BRCA: stage WT n=621, neoGATA3 n=20, OtherMut n=70; PR WT n=625, neoGATA3 n=21, OtherMut n=69). Fisher’s test Chi-square *P<0.05, **P<0.01.

**Supplementary Figure 3:**
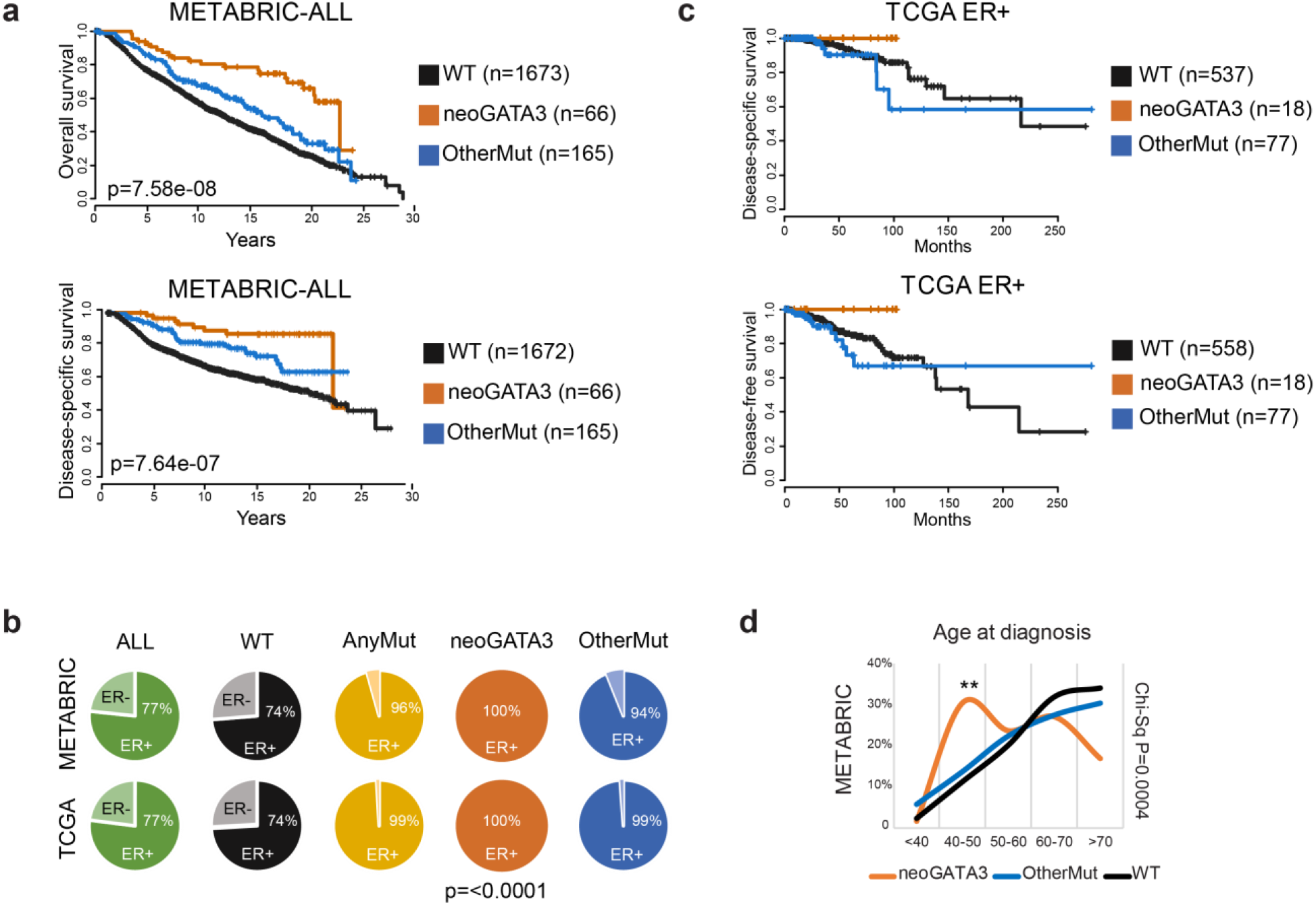
NeoGATA3 mutations are prevalent in ER+ tumors and predict good outcome. (**a**) Kaplan-Meier curves showing overall survival and disease-specific survival of the METABRIC patients stratified according to *GATA3* status. (**b**) Distribution of ER+ or ER-tumors among the indicated subgroups of the METABRIC and the TCGA-BRCA cohorts. AnyMut= any *GATA3* mutation. (**c**) Kaplan-Meier curves showing disease-specific and disease-free survival data of the TCGA ER+ cohort stratified as above. (**d**) Graph showing the age at diagnosis of the METABRIC patients belonging to the three groups (WT n=929, neoGATA3 n=59, OtherMut n=138).

**Supplementary Figure 4:**
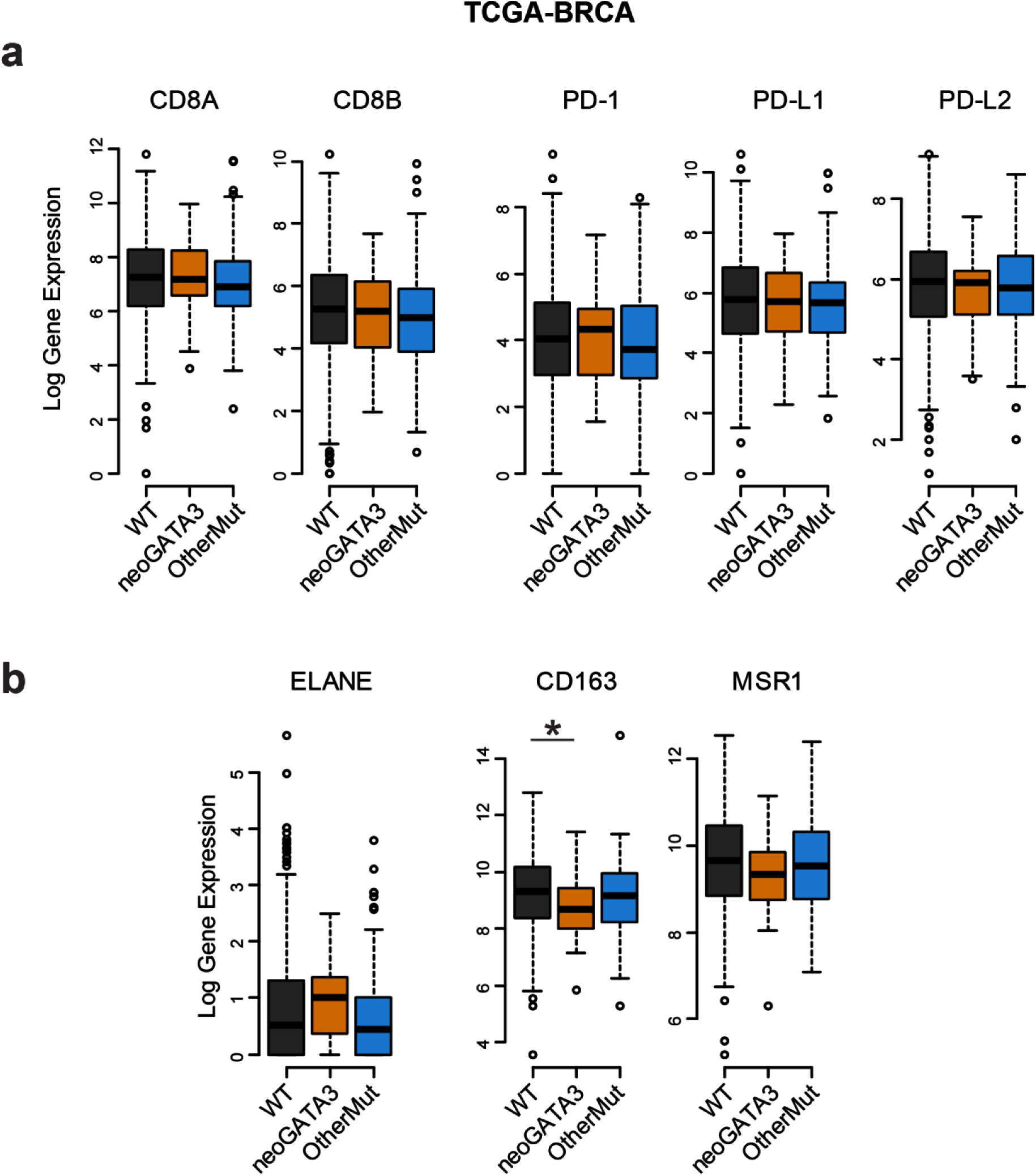
NeoGATA3 mutations are not associated with an immune cell infiltration. (**a,b**) Gene expression levels of the indicated markers of T-lymphocytes, neutrophils, and M2 macrophages in tumors of the TCGA cohort, divided in the three groups according to the *GATA3* status (WT n=609, neoGATA3 n=20, OtherMut n=87). Two-sided Student’s T test *P<0.05.

**Supplementary Figure 5:**
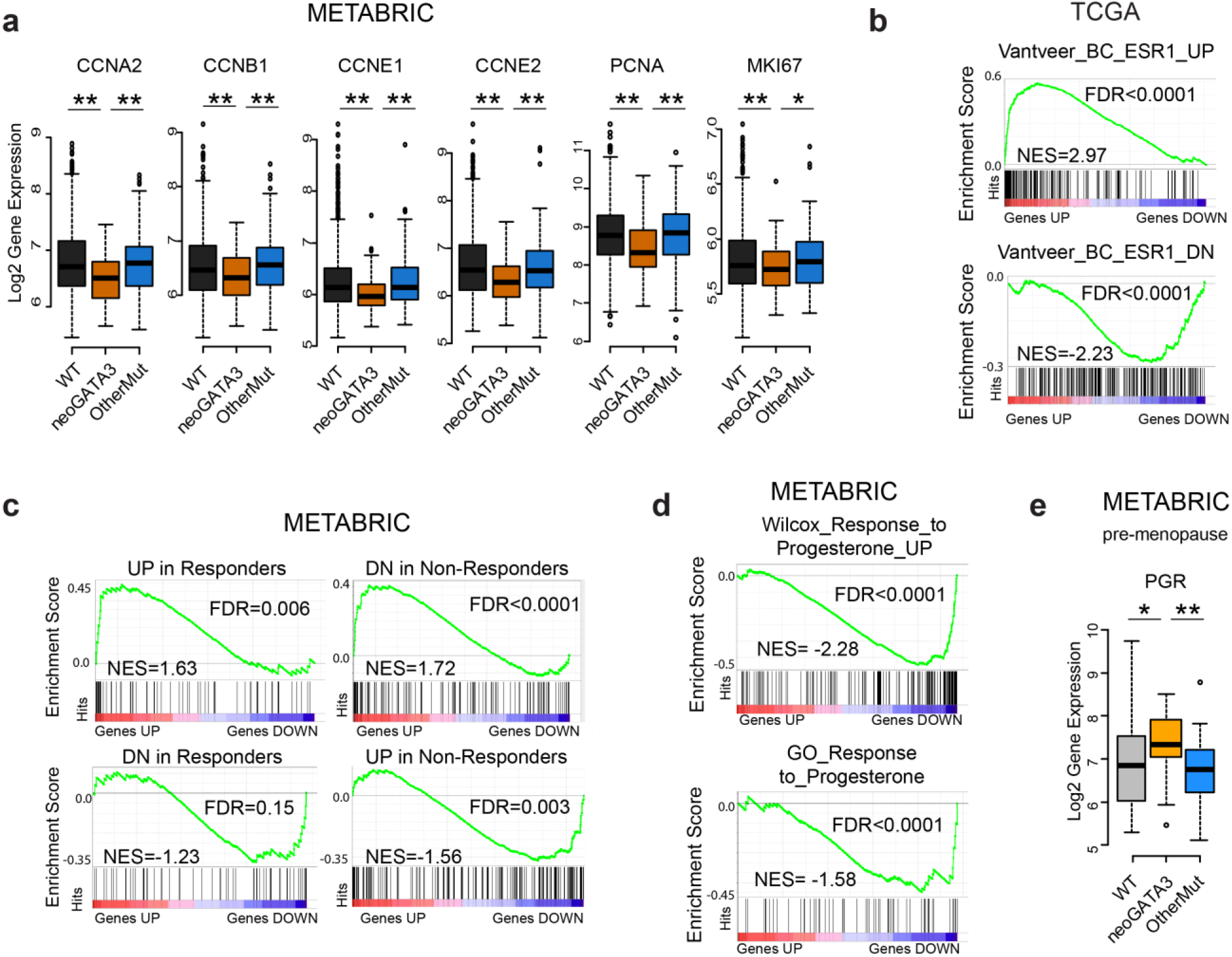
The neoGATA3 protein interferes with the ER-dependent and PR-dependent programs in tumors. (**a**) Gene expression data for the indicated cell cycle-related genes in the METABRIC patients belonging to the three groups (WT n=1189, neoGATA3 n=66, OtherMut n=155). (**b**) Enrichment plots of ER-related signatures within the genes differentially regulated in neoGATA3 tumors versus all others from the TCGA cohort. (**c**) Enrichment plots for genesets defined by Ross-Innes et al. comparing gene expression in tumors responding to endocrine therapy and tumors with poor response. GSEA was performed on genes differentially regulated in the neoGATA3 tumors from the METABRIC cohort. (**d**) Enrichment plots for two progesterone-related genesets among the differentially expressed genes in the METABRIC neoGATA3 patients compared to all other METABRIC ER+. (**e**) Gene expression data for the *PGR* gene in pre-menopausal METABRIC ER+ patients of the three groups (WT n=157, neoGATA3 n=21, OtherMut n=30). Two-sided Student’s T test *P<0.05, **P<0.01

**Supplementary Figure 6:**
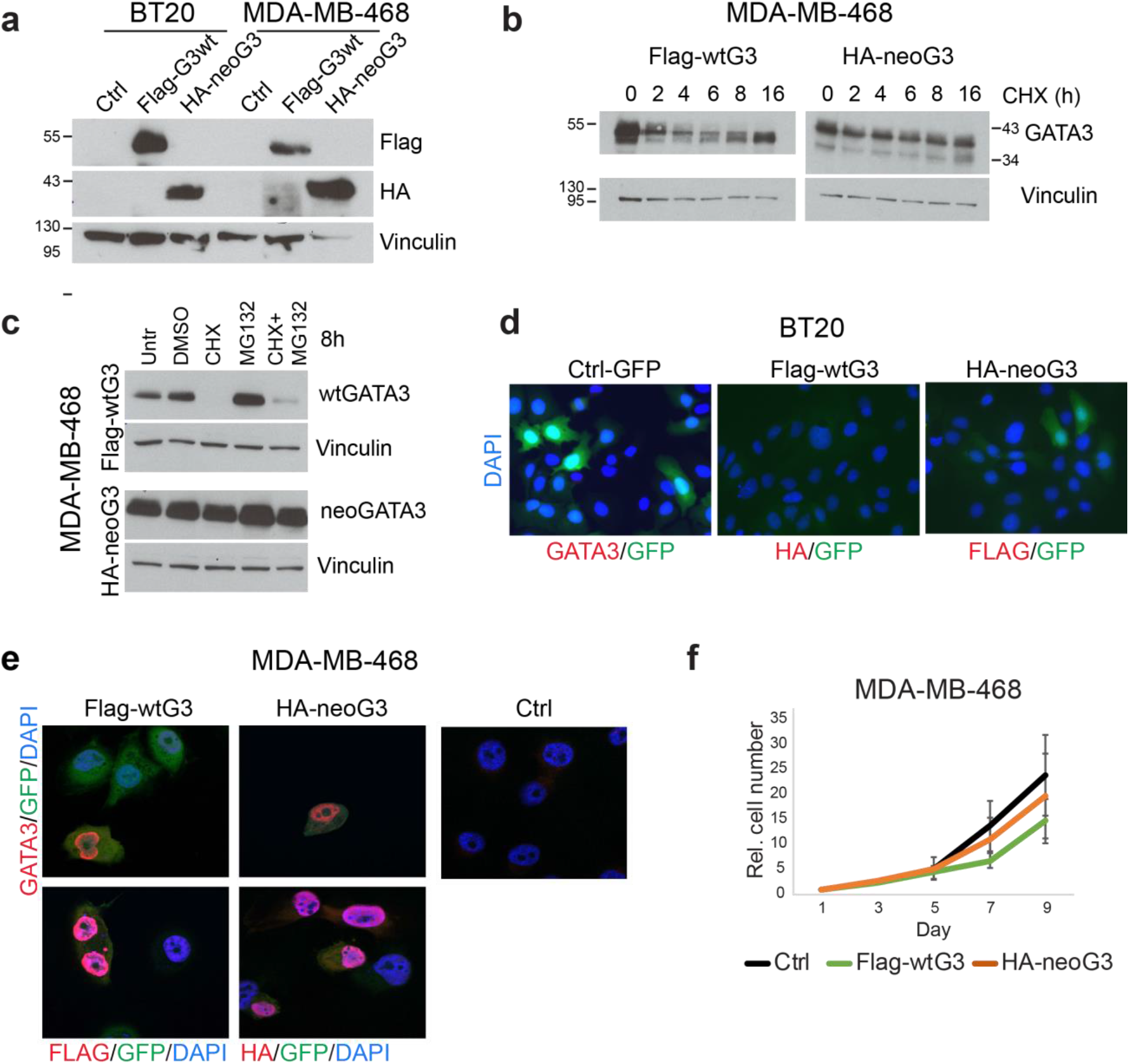
Expression of neoGATA3 in GATA3-negative BC cells. (**a**) Western blot showing the detection of the Flag and HA tags in BT20 and MDA-MB-468 cells transduced with Flag-wtGATA3 (Flag-wtG3) or HA-neoGATA3 (HA-neoG3). Vinculin was used as loading control. (**b**) Western blot showing the protein level of wtGATA3 or neoGATA3 expressed in the GATA3-negative MDA-MB-468 BC cells, after treatment with cycloheximide (CHX) for the indicated time. Vinculin was used as loading control. (**c**) Western blot showing expression of wtGATA3 or neoGATA3 in MDA-MB-468 cells transduced with either wtGATA3 (top) or neoGATA3 (bottom) after treatment with CHX, MG132, or both. Vinculin was used as loading control. (**d**) Representative images showing the negative controls of the experiment shown in Figure 3B. Ctrl-transduced BT20 cells are GATA3-negative, Flag-wtG3-transduced cells are HA-negative, HA-neoG3-transduced cells are Flag-negative. DAPI was used to counterstain nuclei, GFP was expressed by the lentiviral vector used for the transduction. (**e**) Immunofluorescence using the GATA3 antibody (top panels) or tag-specific antibodies (bottom panels, left: Flag, right: HA) in MDA-MB-468 cells expressing either Flag-wtG3 or HA-neoG3, or Ctrl-transduced cells, as indicated. DAPI was used to counterstain nuclei, GFP was expressed by the lentiviral vector used for the transduction. (**f**) Growth curve of MDA-MB-468 cells transduced with the indicated constructs. Data are represented as mean ± standard deviation of at least three independent experiments.

**Supplementary Figure 7:**
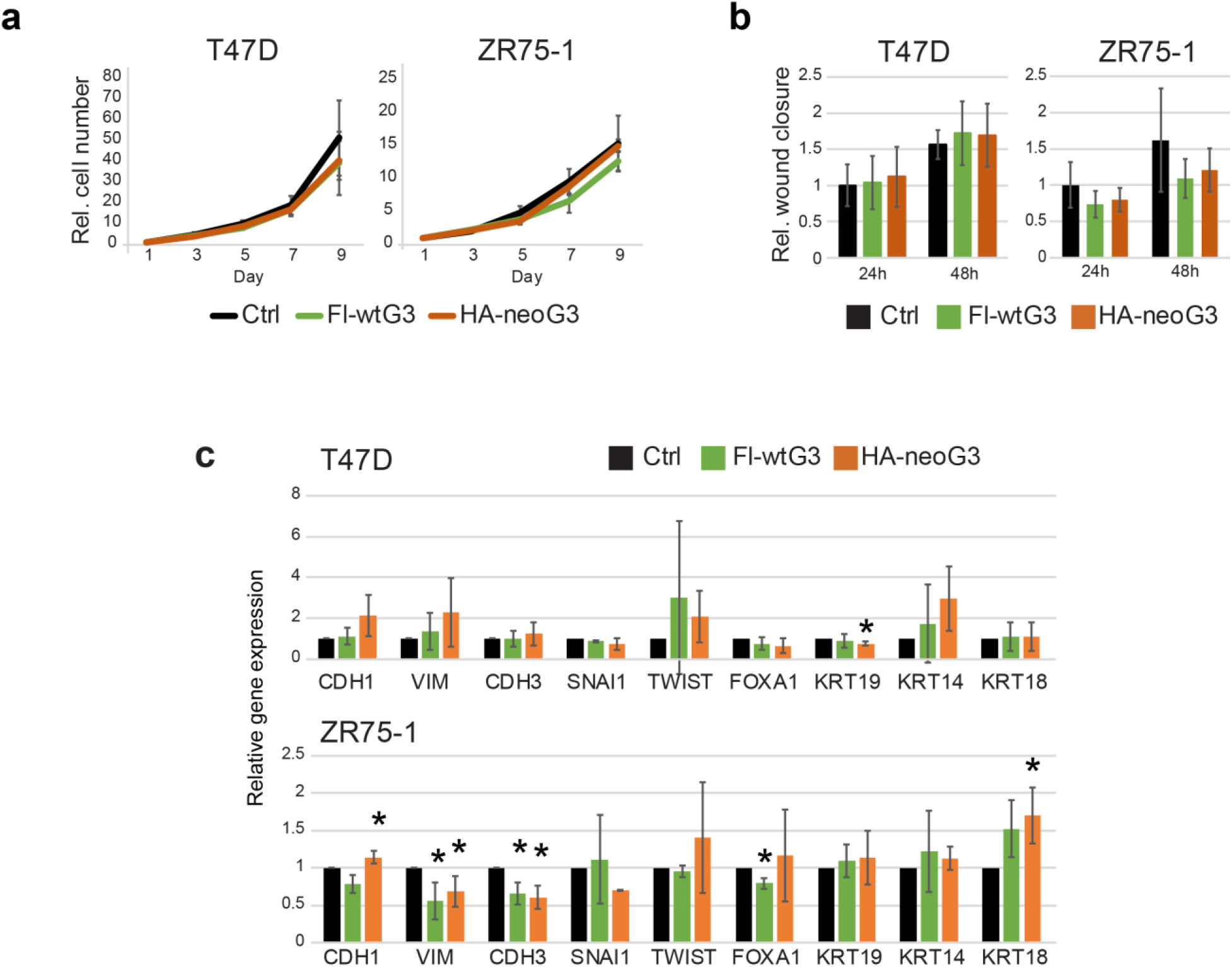
Expression of wtGATA3 and neoGATA3 in luminal GATA3-positive BC cells. (**a**) Growth curves of T47D and ZR75-1 cells transduced with the indicated constructs. Data are represented as mean ± standard deviation of at least three independent experiments. (**b**) Graphs showing the relative wound closure in a scratch assay performed with T47D and ZR75-1 cells transduced with the indicated constructs after 24h or 48h. Data are represented as mean ± standard deviation of at least three independent experiments. (**c**) Graphs showing the relative expression levels of differentiation-related genes in T47D and ZR75-1 cells transduced as indicated. All values are normalized to Ctrl-transduced cells. Data are represented as mean ± standard deviation of at least three independent experiments. Two-sided Student’s T test *P<0.05.

**Supplementary Figure 8:**
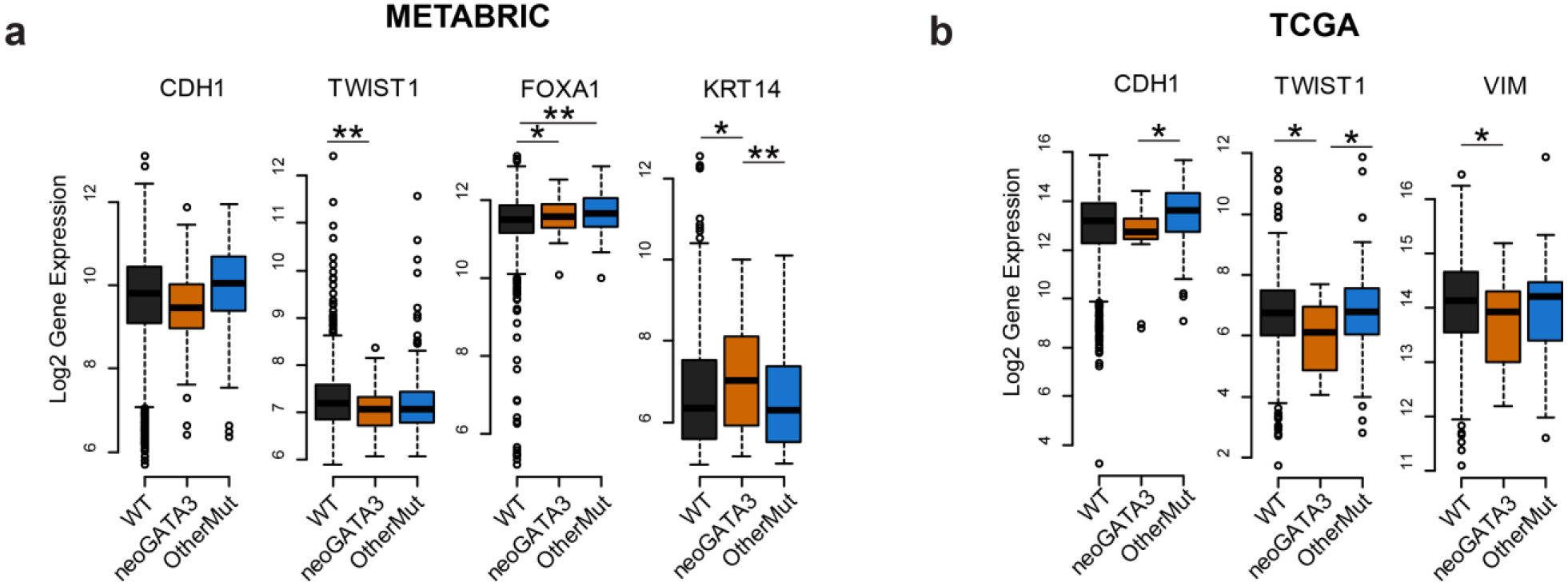
NeoGATA3 tumors express lower levels of some EMT genes. (**a,b**) Gene expression data from the METABRIC (**a**) and from the TCGA cohort (**b**), showing the levels of EMT markers in patients of the three groups. *P<0.05, **P<0.01

**Supplementary Figure 9:**
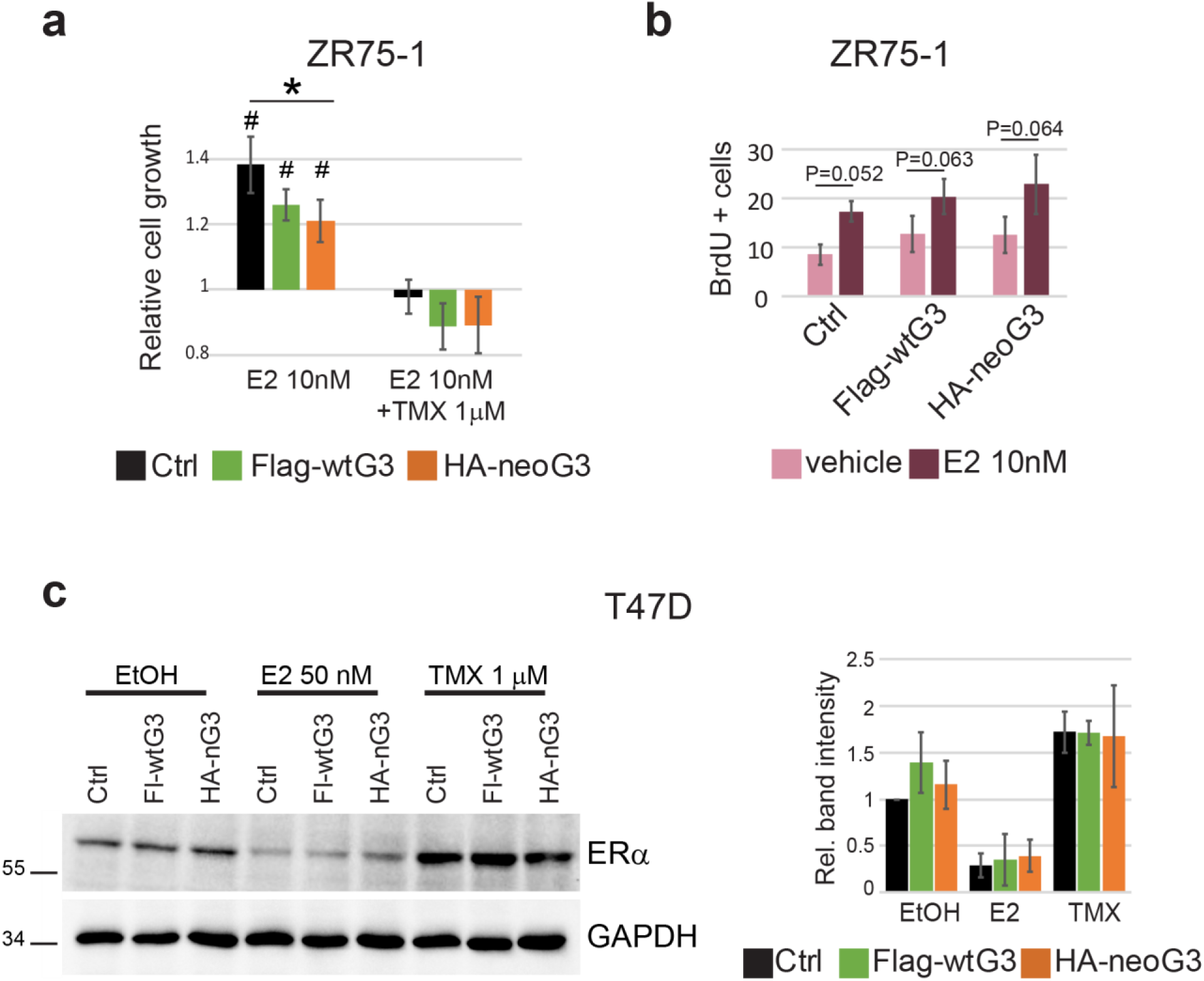
NeoGATA3 interferes with the ER-dependent program *in vitro*. (**a**) Graph showing the relative cell viability of ZR75-1 cells transduced with the indicated constructs and treated with E2 alone (10nM) or in combination with TMX (1μM) for 72h after 48h in HD medium. (**b**) Graph showing the percentage of BrdU+ cells in ZR75-1 cells treated with E2 (10nM) for 24h. Data are shown as mean ± standard deviation of at least three independent experiments. *P<0.05, **P<0.01, #P<0.05 compared to the vehicle control. (**c**) Western blot showing ER expression in T47D cells transduced with the indicated constructs and treated 24h with vehicle (EtOH), E2 (50nM), or TMX (1μM) after 48h in HD medium. GAPDH was used as loading control. Band intensity relative to Ctrl cells treated with vehicle was quantified in three independent experiments and the results are shown on the right as mean ± standard deviation.

